# Combined Dendritic Cell And Anti-TIGIT Immunotherapy Potentiate Trail+ Memory NK Cells Against HIV-1 Infected Cells

**DOI:** 10.1101/2024.04.09.587160

**Authors:** I Sánchez-Cerrillo, O Popova, M Agudo-Lera, I Tsukalov, M Calvet-Mirabent, I de los Santos, L García-Fraile, P Fuentes, C Delgado-Arévalo, J Alcain, N Sánchez-Gaona, M Lázaro-Díez, C Muñoz-Calleja, Arantzazu Alfranca, M Genescà, JG Prado, Vladimir Vbrnac, Alejandro Balazs, MJ Buzón, M.L Toribio, MA Muñoz-Fernández, F Sánchez-Madrid, E Martín-Gayo

**Author notes:** Corresponding author: Enrique Martin-Gayo Ph.D.; Assistant Professor, Universidad Autónoma de Madrid Immunology Unit, Hospital de la Princesa, Calle de Diego de León, 62, 28006 Madrid, Spain.

## Abstract

Natural Killer (NK) cells are promising tools for the development of immunotherapies targeting persistently infected CD4+ T cells to potentially achieve remission in people with HIV-1 (PWH). However, the chronicity of HIV-1 infection limits the functional properties of NK cells, and additional approaches are needed to potentiate their cytotoxic activity against HIV-1-infected cells. In the present study, we analyzed the reinvigoration of functional NK cells from PWH after priming with autologous dendritic cells (DC) stimulated with nanoparticles containing Poly I:C (Nano-PIC). We show that improved natural cytotoxic function in NK cell from PWH associates with increased proportions of NKG2C+CD57- precursors of memory NK, which eliminate HIV-1 infected CD4+ T cells mainly through the TRAIL receptor. In addition, expression of TIGIT but not TIM3 limited increase in NKG2C+ memory NK cell precursors and associated with persistent dysfunctionality of NK cells after stimulation with Nano PIC-DC. Blockade of TIGIT restored functional capacities of NK cell from PWH eliminating HIV-1 infected cells *in vitro*. Moreover, combining of NK cell and Nano-PIC-DC with anti-TIGIT mAbs immunotherapy limited the expansion of HIV-1 infected cells in humanized immunodeficient NSG mice transplanted with CD4+ T cells from PWH *in vivo*. Such viral control was associated with preserved NKG2C memory NK cell precursors, increased expression of granzyme B and TRAIL on NK in tissue from transplanted NSG mice. Together, combination of Nano-PIC DC and anti-TIGIT antibodies may be a promising strategy to increase the efficacy of immunotherapies aimed at HIV-1 cure.

**One sentence summary:** Stimulation of memory NK with a combination of DC and anti-TIGIT antibodies increase their ability to eliminate HIV-1 infected CD4+ T cells *in vitro* and *in vivo*.

## INTRODUCTION

Antiretroviral treatment (ART) is very effective inducing almost complete viral suppression and reduction to undetectable plasma viral load in people living with HIV-1 (PWH). However, this treatment does not eliminate persistent HIV-1 reservoirs in latently infected CD4+ T cells (*1–3*), which are present in different tissues such as lymphoid organs and gastrointestinal mucosa (*3–7*). A fraction of these latently infected T cells contains complete viral sequences and can produce replication-competent HIV-1 particles ((*8, 9*). It has been proposed that the proliferation of these latently infected cells allows the maintenance of persistent HIV-1 reservoirs (*10–12*). In addition, the chronicity of the persistent HIV infection and the sporadic reactivation of latently infected cells in PWH leads to residual inflammation and continuous stimulation of immune cells with antiviral capacities such as CD8+ T cells and Natural Killer (NK) cell. Such hyperstimulation eventually leads to an exhausted state in these immune cells that is characterized by the expression of multiple inhibitory checkpoint receptors, reduced functional properties and impaired ability to eliminate infected cells (*13, 14*). In some cases, ART does not seem to completely restore such a dysfunctional, exhausted state in immune cells and longer periods of treatment may be required to recover responsiveness in these cells (*15–18*). Thus, new approaches are needed to develop a more effective antiviral immune response against HIV-1 and to achieve a reduction of persistent viral reservoirs and potentially, a cure. Several studies have previously focused on enhancing HIV-specific T cell responses ((*19–25*) from PWH on ART, including the stimulation with activated dendritic cells (DCs) loaded with HIV-1 peptides (*19, 26–29*) and/or the blockade of checkpoint receptors (*30, 31*). While these approaches have improved T-cell mediated targeting of HIV-1 infected cells *in vitro* and *in vivo*, they may not be sufficient to reduce viral reservoirs to a level capable of promoting spontaneous control or remission of infection.

NK cells are innate immune cells with antiviral capacities that have been shown to be essential to control HIV replication *in vivo* in animal models (*32–38*) and in spontaneous and post-treatment controllers (*15, 39–41*). Although some studies point to their promising therapeutic potential against HIV-1, modulation of NK cells has been less explored (*37, 42–47*). Therefore, enhancing the activity of these cells has currently become an area of active research. NK cells are a heterogeneous linage of innate immune cells that are capable of detecting and eliminating virus-infected cells either by directly recognizing ligands for activating receptors or antibodies bound to viral proteins expressed by target cells. These two killing mechanisms are known as natural and Antibody-Dependent Cellular (ADCC) Cytotoxicity (*48–53*), and are dependent on the expression of a repertoire of activating receptors such as NKG2D or NKG2C and CD16 expressed by NK cells at different stages of maturation, respectively. In this regard, NKG2C+ cells are a subset included in CD56dim CD16+ and CD56lo/- CD16+ NK cells initially identified as an expanded population in CMV infected individuals that is characterized by a primed state, including higher cytotoxic potential and enriched ADCC activity, and therefore, this subset has been considered to have memory-like properties (*54, 55*). These cells originate from NKG2C-NKG2D+ precursors giving rise to immature NKG2C+ CD57- precursors and finally, mature NKG2C+ CD57+ memory cells (*56–59*). In the case of HIV-1 infection, memory NKG2C+ NK cells have been shown to be capable of recognizing HLA-E expressing infected CD4+ T cells loaded with viral peptides (*60, 61*). In addition, NKG2C+ memory NK cells were also associated with remission and functional cure of HIV infection (*62*). Thus, inducing and potentiating memory NKG2C+ NK cells may be beneficial to improve immunotherapy against HIV-1 and this possibility has not been explored.

Among different tools useful to improve antiviral immune responses, DC are myeloid innate immune cells that are able not only to present antigens and mediate polarization of T cells (*63, 64*) but they also have the ability to support maturation and activation of NK cells (*65–67*). Recently, we observed that the activation of intracellular innate RNA sensors such as RIG-I increases expression of ligands for NKG2D and the ability of DC to activate cytotoxic NK cells in the autoimmune disorder Sjogreńs Syndrome ((*68*). Moreover, DC modified by agonists of the STING and TLR3 pathways can be used as an immunotherapy to restore HIV-1 specific CD8+ T cell responses from PWH (*19*). However, the potential of DC to enhance cytotoxic function of NK cells from PWH and those clinical and immunological parameters associate with increased NK functional response have not been studied in detail. In this regard, the use of nanoparticles is an attractive choice to facilitate the selective targeting of intracellular sensors such as RIG-I, which may be beneficial to enhance NK cell-activating function in DC and to improve immunotherapies against HIV-1.

Here, we show that nanoparticles loaded with Poly I:C increase the ability of DC from PWH to activate autologous NK cells and to induce functional restoration and increased cytotoxic activity of these cells against HIV-1 infected CD4+ T cells. Such enhancement of NK cell activity was associated with increased proportions of memory-like NKG2C+ TRAIL+ NK cells. Moreover, functional restoration of NK was negatively associated with TIGIT expression and positively correlated with TIM3. Finally, blockade of TIGIT was capable of restoring cytotoxic activity of dysfunctional NK cells against p24+ cells *in vitro*. Moreover, it was associated with reduced expansion of infected cells and with an enrichment in NKG2C+ TRAIL+ NK cells in lymphoid tissue *in vivo* in humanized mice. Together, our study indicates that combined blockade of TIGIT and stimulation with Nano-PIC DC immunotherapy can potentially be used as a strategy to improve NK cell cytotoxic functionality and to reduce persistent HIV-1 replication.

## MATERIALS AND METHODS

### Study Participants

We recruited n=28 PWH on ART for a median of 9.2 years (min-max; 1,5-25) and characterized by undetectable plasma viremia (<20 copies HIV-1 RNA/mL), CD4+ T cell count higher than 500 cell/ul (median 835; min-max, 596-1639) and no co-infection with hepatitis C virus for the study from the Infectious Diseases Unit from Hospital Universitario La Princesa. Median age was 52 (min-max, 25-75) and 96% of participants were male. Additional clinical characteristics and treatment regimens are summarized in Supplementary Table 1.

### Generation of DC and stimulation with Poly I:C loaded nanoparticles

DC were generated *in vitro* from circulating adherent Mo from HIV negative donors and from PWH in the presence of recombinant human 100 IU/mL GM-CSF and 100 IU/mL IL-4 (Preprotech) for 6-7 days. Subsequently, DCs were cultured for additional 16h in the presence of either 1μg/ml soluble Poly I:C (Invivogen) (s-PIC) or complexes of polymeric nanoparticles (TRANS-ITX2-Mirus bionova) containing the same concentration of Poly I:C (Nano-PIC). Controls of DC cultured in the presence of just media or empty nanoparticles, respectively. In these experiments, phenotypical analysis of expression of cytokines, maturation markers and ligands for NK receptors on DC cultured under the described conditions was carried out by FACS (see Flow cytometry section).

### Natural cytotoxicity and Antibody-dependent cellular cytotoxicity in vitro assays

To address killing abilities of NK cells stimulated with Nano-PIC DC we used two different assays with target cell lines. To address natural cytotoxic function, we co-cultured NK cells with the MHC-devoid target cell line K562 transfected with GFP (NIH AIDS reagent program Ref 11699). Specific killing was determined by flow cytometry quantifying loss of GFP expression and acquisition of a viability dye. For Antibody-dependent cellular cytotoxicity (ADCC) we co-cultured NK cells in the presence of the cell line CHO cell line transfected with HIV-1 gp120 selected as specified by the provider (NIH AIDS reagent program Ref 2239) alone or preincubated with a cocktail of three HIV-1 specific broadly neutralizing antibodies (10μg/mL VCR01; 5μg/mL PGT121; 1μg/mL 3BNC117). Cell death was quantified by acquired staining viability dye staining. ADCC activity was calculated after the sustraction of the basal killing of the CHO target cell line in the absence of bNAbs.

### Immunomagnetic selection of NK, co-culture with NanoPIC-DC and functional assays CD4+ T cells from PWH

NK cells were isolated by negative immunomagnetic selection (STEM cell) from the blood of HIV negative donors and aviremic PWH on ART (purity >90%) and co-cultured with autologous DC pre-stimulated with either sPIC or Nano-PIC or their control conditions in the presence of 50 IU/mL IL-2 (Prepotech). In some experiments, levels of intracellular expression of IFNγ, TNFα, granzyme B and CD107a was addressed in NK from PWH cultured in the mentioned conditions. Subsequently, primed NK cells from PWH were cultured with autologous CD4+ T cells also isolated by negative immunomagnetic selection (DynaBeads Human CD4+ T cell isolation kit, Life technologies) in the presence of 50nM Romidepsin (Selleck Chemicals) to induce viral reactivation and 30μM Raltegavir (Selleck Chemicals) to prevent viral spread. After 16h, proportions of p24+ HIV-1 infected CD4+ T cells was addressed by Flow cytometry. In some of these assays, expression of CD56 vs CD16, NKG2C and CD57 defining different subsets of memory and effector NK cells, checkpoint receptors (TIGIT, PD1 and TIM3) and TRAIL within CD56dim CD16+ and CD56lo/- CD16+ populations were tested and associated with functional ability to kill HIV-1 infected CD4+ T cells. In some experiments, expression of the TRAIL ligand DR4 (BioLegend), TIGIT ligand CD155/PVR (BioLegend) and NKG2D/C ligands MICa/b or HLA-E, respectively, was also tested in p24- vs p24+ isolated CD4+ T cells from PWH after reactivation with PHA + IL-2. Finally, impact of anti-Trail (RyD Systems), anti-NKG2C (RyD Systems), anti-NKG2D (RyD Systems) and anti-TIM3 (RyD Systems) and anti-TIGIT (RyD Systems), or the corresponding mouse IgG isotypic mAb (Biolegend) and goat IgG (SIGMA) was assessed in the mentioned functional NK cell-mediated killing of autologous HIV-1 infected CD4+ T cells from PWH described above (Supplemental Table 2).

### Flow cytometry

Maturation of DC was addressed using anti-CD86 and either intracellular staining with anti- IFNβ (Pbl assay science), anti-IL-12 (RyD Systems) in the presence of 0.25 μg/mL Brefeldin A and 0.0025 mM Monensin (SIGMA) or in combination with anti-MICa/b, anti-ULBP1 or anti-HLA-E from Biolegend. (Supplemental Table2).

The analysis of NK cell subpopulations was performed using a Ghost Dye Red 780 viability dye (TONBO Biosciences), anti-CD56, anti-CD16, in combination with either IFNγ, CD107a, Granzyme B, TNFα mAbs (BioLegend) in presence of PMA and Ionomycin for 1 hour and Brefeldin A and Monensin after 4 hours or NKG2D, NKG2A, NKG2C, CD57, TIGIT, TIM3, PD- 1 or TRAIL mAbs (BioLegend) (Supplemental Table2).

In functional assays proportion of infected CD4+ T cells was address by intracellular staining of p24 (KC57, Beckman Coulter) in combination with anti-CD3, anti-CD4 (BioLegend) antibodies and in the presence of the mentioned viability dye. Samples were analyzed in a BD Canto II and Lyric instruments. Analysis of flow cytometry data was performed using FlowJo v10.6 software (Tree Star).

### Analysis of memory NK cell and impact of combined Nano-PIC DC anti-TIGIT immunotherapy in humanized mice in vivo

A total of n=13 NOD/SCID IL2R-y-/- (NSG) mice transplanted with human BM, fetal liver, and thymus (BLT-mouse) and fetal CD34+ HSCs were generated at the Human Immune System Core from the Ragon Institute and Massachusetts General Hospital in collaboration with Dr. Vladimir Vrbanac and Dr. Alejandro Balazs as previously described (*69*). Mice were housed in ventilated racks and fed autoclaved food and water at a pathogen-free facility. Intraperitoneal injections of 2.5μg/0.02kg rhIL-15 (R&D systems; Supplemental Table 2) were performed for 3 weeks to increase reconstitution and maturation of human NK cells in hBLT mice. Human immune reconstitution was monitored for 17 weeks prior to infection with HIV-1 and optimal reconstitution was considered with proportions of human CD45+ lymphocytes over 25%. Viral stocks were generated by transfection of HEK-293 (ATCC® CRL-1573) cells with the plasmid “HIV-1 JR-CSF Infectious Molecular Clone (pYK-JRCSF)” (National Institutes of Health [NIH] AIDS Reagent Program, catalog number 2708) using the X-tremeGENE 9 DNA Transfection Reagent (Roche) according to manufacturer’s instructions. 72 h after transfection, supernatants were collected, filtered, concentrated and resuspended in 1x PBS. The JRCFS viral stock was stored at −80°C for further experiments. The virus TCID50 was determined in TZM-bl cells (NIH AIDS Reagent Program, catalog number 8129) by limiting dilution using the Reed and Muench method, as previously described. Dr. Julia García Pradós group at IRSICaixa, Barcelona, Spain produced and provided the viral stocks for the study. Reconstituted hBLT mice were either intravenously infected (n=8) with 5,000 TCID50 of JRCSF or left uninfected (n=5). Body weight, HIV-1 plasma viral load, CD4+ T cell counts and proportions of memory and effector NK cell subsets in the blood were monitored during 3 weeks.

To address the efficacy on anti-TIGIT modulation of NK cell-Nano-PIC DC immunotherapy eliminating HIV-1 infected CD4+ T cells from PWH in vivo, we intravenously transplanted n=27 immunodeficient NOD/SCID IL2R-y-/- (NSG) mice with either CD4+ T cells from PWH (n=3 different donors in three independent experiments) NK cells, Nano-PIC DC from the same PWH at a 1:2:4 (DC:NK:CD4) ratio. Mice receiving Nano-PIC DC immunotherapy were split into two groups that were administered intraperitoneally every 48h with either humanized anti-TIGIT (Tiragolumab, Selleck Chemical) (n=14 from three independent experiments) or humanized IgG1 isotypic (BioXCell) (n= 13 from three independent experiments) mAbs and n=10 mice were included on each group in two independent experiments (Supplemental Table 2). As control group, NSG mice were also transplanted only with CD4+ T cells from the same PWH (n=12, from three independent experiments). Proportions of p24+ cells in human CD45+ CD4+ T cells in the peripheral blood, liver and spleen at 7 days post-injection analyzed by flow cytometry.

### RT-qPCR quantification of Plasma HIV-1 viral load in humanized mouse models

HIV RNA was extracted from plasma samples collected from either HIV-1 infected or uninfected humanized hNSG mice transplanted with CD4+ T cells from PWH in the absence or presence of immunotherapy using Nucleospin^TM^ RNA Virus kit (Cultek, Macherey-Nagel) following the manufacturer’s instructions. Reverse transcription of RNA to cDNA was performed with SuperScript^TM^ III Kit (Invitrogen) in accordance with the instructions provided by the manufacturer, and cDNA was quantified by quantitative PCR (qPCR) using primers and probes specific for the 1-LTR region (LTR forward 5′-TTAAGCCTCAATAAAGCTTGCC-3′ and LTR reverse 5′-GTTCGGGCGCCACTGCTAG-3′; LTR probe 5′- CCAGAGTCACACACCAGACGGGCA-3′), and TaqMan^TM^ Gene Expression Master Mix (Thermo Fisher). Samples were analyzed in an Applied Biosystems QuantStudio5 system. Quantification of DNA was performed using a standard curve, and values were normalized to 1 million CD4+ T cells.

### Histological analysis of tissue sections from humanized mice

Spleens were collected from hBLT mice and NSG transplanted mice and paraffin-embedded and segmented in fragments of 2 μm of thickness in a Leica microtome. Tissue sections deparaffinized, hydrated, and target retrieval were performed with a PT-LINK (Dako) previous to staining.

For paraffin-preserved tissue, rat anti-human Granzyme B (eBioscience), mouse anti-HIV-1 P24 (Dako), goat anti-human TRAIL (RyD Systems) and rabbit anti-human NKG2C (Abcam) as primary antibodies; and goat anti-rabbit AF488 (Invitrogen), donkey anti-rat AF594 (Jackson ImmunoResearch), donkey anti-goat AF568 (Invitrogen) and donkey anti-mouse AF647 (ThermoFisher) as secondary antibodies (Supplemental Table 2). Images were taken with a Leica TCS SP5 confocal and processed with the LAS AF software. Granzyme B+ and HIV-1 p24+ cell counts and colocalization and distribution of some of these markers were analyzed with ImageJ software.

### Statistical analysis

Statistical significance of differences between cells from different or within the same patient cohort under different treatments were assessed using Mann Whitney U or Wilcoxon matched-pairs signed-rank tests. Chi-square with Yate’s correction was used to compare differences in proportions of some parameters within different cell/subject populations.

Non-parametric Spearman correlations were used to test both individual correlations and to generate a correlation network. Statistical analyses were performed using GraphPad Prism 9.0 software.

### Ethic statement

All subjects participating in the study gave written informed consent, and the study was approved by the Institutional Review Board of Hospital Universitario de La Princesa (Protocol Registration Number 3518) and following the Helsinkís declaration. For *in vivo* experiments, mice were housed at the animal facility from Centro de Biología Molecular Severo Ochoa (CBM) in accordance with the institution’s animal care standards. Animal experiments were reviewed and approved by the local ethics committee and were in agreement with the EU Directive 86/609/EEC, Recommendation 2007/526/EC and Real Decreto 53/2013.

## RESULTS

### Nanoparticle stimulation of intracellular RNA sensors increase ability of DC to activate cytotoxic CD16+ NK cells

To determine whether stimulation of DCs with soluble (Sol-PIC) or nanoparticle-encapsulated Poly I:C (Nano-PIC) may differentially regulate functional maturation in these cells and their ability to stimulate NK cells, we first analyzed cell viability, expression of different cytokines and induction of maturation markers after *in vitro* stimulation (Supplemental Figure 1A). Treatment of DC with Nano-PIC induced a mild but significant reduction of cell viability compared to cells treated with Sol-PIC, although it remained over 60% (Supplemental Figure 1B, left panel). As shown in Figure 1A, while both stimulation methods induced IL-12 and IFNβ secretion by DC, Nano-PIC more significantly increased intracellular expression of these cytokines at 6 and 8h compared to cells treated with Sol-PIC (Figure 1A). Likewise, the expression of costimulatory molecules such as CD86 was also more efficiently upregulated at 6, 8hr and 16h after stimulation in DC primed with Nano-PIC (Supplemental Figure 1B, right panel; Figure 1B). Interestingly, opposite to cells exposed to Sol-PIC, DC primed with Nano-PIC more efficiently induced significant higher expression of ligands for the activating NK receptor NKG2D such as MICa/b and ULBP1 as well as HLA-E, which are the ligands for both the activating and inhibitory receptors NKG2C/D and NKG2A, respectively (Supplemental figure 1C; Figure 1C upper panel). The same effects were also observed in DC from PWH exposed to Nano-PIC (Figure 1C, lower panel). The expression of these ligands was more enriched on activated CD86+ DCs after Nano- PIC stimulation (Supplemental Figure 1D). Therefore, these data indicate that Nano-PIC DC may display increased abilities to activate NK cells. To address this possibility, we compared the ability of DC treated with empty nanoparticles or Nano-PIC to increase maturation and activation of autologous NK cells *in vitro*. To this end, expression of IFNγ and CD107a on CD56dim CD16+ cells was analyzed (Supplemental Figure 1E). An increase in the expression of the degranulation marker CD107a (Figure 1D, upper panel) and proportions of CD107a+ IFNγ+ (Figure 1D, lower panel) cells within the CD16+ CD56dim NK subset were observed in response to Nano-PIC DC. Notably, this effect was dependent on cell-to-cell contact of NK cells with Nano-PIC DC in a transwell assay (Figure 1E). Also, expression of NKG2D and a trend to higher NKG2C expression, was observed in CD16+ CD56dim and CD56lo/- cell populations present in pre-isolated NK cells cultured with Nano-PIC DC (Supplemental Figure 2A-B-C).

**Figure 1.**
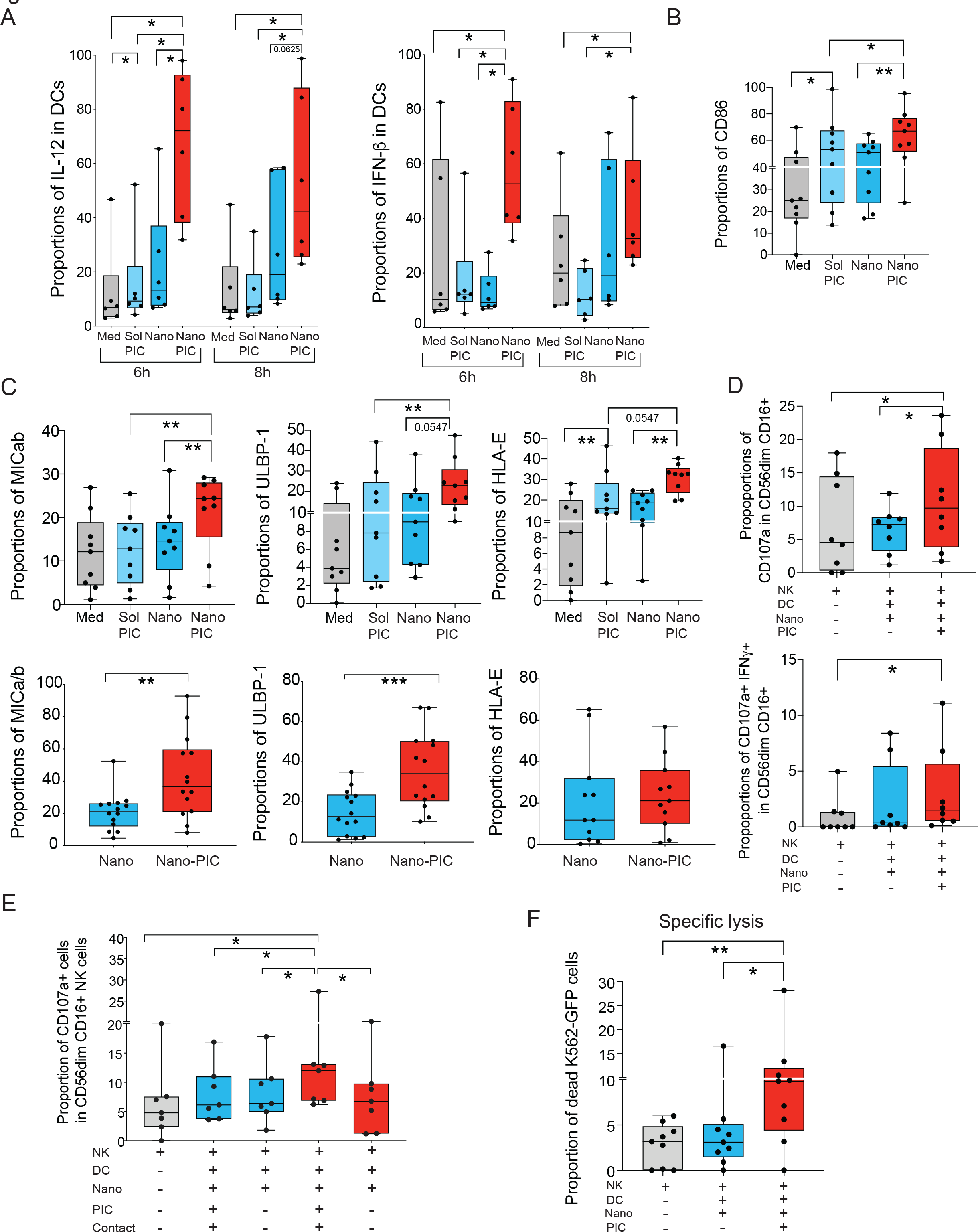
Analysis of cytokine production, maturation and expression of ligands for NK receptor in DC stimulated with nanoparticle-encapsulated Poly I:C. (A-B-C): Intracellular expression of IL-12 and IFNβ at 6 and 8h in n=6 independent experiments (A) or surface levels of CD86 (B) and NK receptor ligands MICa/b, ULBP-1 and HLA-E (C, upper panel) after 16h of stimulation with either soluble (Sol) or Nanoparticle-encapsulated (Nano) Poly I:C (PIC) in n=8 independent experiments with healthy donors (HD). Expression of NK receptor ligands on Nano- PIC DCs from PWH is also shown in the lower panel in C. (D): Analysis of proportions of total CD107a+ (upper plot) and IFNγ+CD107a+ cells (lower panel) on CD56dim CD16+ NK cells cultured in the absence or the presence of DC stimulated either with empty or PIC-loaded nanoparticles in the same culture (n=8 independent experiments), Proportions of CD107a+ CD56dim CD16+ NK in the absence or the presence of Nano-PIC DC with or without cell contact a transwell assay (E). (F): Proportions of dead target K562-GFP cell line after culture in the presence of NK ells alone or co-cultured with DC treated with empty or PIC-loaded nanoparticles (n=9). Statistical significance was calculated using a two tailed Wilcoxon pair matched test. *p<0.05; **p<0.01.

To address whether augmented CD107a expression reflected an increase in the cytotoxic capacities of NK cells stimulated with Nano-PIC DC, we performed killing assays in the presence of the target cell line K562 transfected with GFP (Supplemental figure 2D). As shown in Figure 1F, NK cells primed with Nano-PIC DC were able to kill significantly higher proportions of K562- GFP cells, compared to untreated NK or cells exposed to DCs primed with empty nanoparticles. In addition, a tendency to a higher ADCC activity against a HIV1-gp120-expressing target cell line precoated with a mix of HIV-1 specific bNAbs was observed in the presence of NK cells stimulated with Nano-PIC DC (Supplemental Figure 2E). Together, the data suggest that nanoparticle-loaded Poly I:C efficiently improves the ability of DC to support the activation of functional cytotoxic NK cells.

### Cytotoxic activity of NK cells from PWH stimulated with Nano-PIC DC associates with the induction of memory NKG2C+ cells

We next assessed whether NK cells from PWH could more efficiently eliminate autologous HIV- 1 infected CD4+ T cells after stimulation with Nano-PIC DCs. To this end, we tested the ability of NK cells from our cohort of aviremic PWH for at least 1 year on ART (Supplemental Table 1) to eliminate autologous p24+ HIV-1 infected CD4+ T cells *in vitro* in the presence of the latency reversal agent (LRA) romidepsin and the antiretroviral drug raltegravir (Supplemental Figure 3A). When analyzing functional assay results considering cells from all participants tested, we did not observe any significant differences in proportions of p24+ CD4+ T cells in culture in the presence of NK cells treated with Nano-PI DC, Nano-DC or just media compared to baseline levels (Supplemental Figure 3B). However, we were able to identify two distinct groups of PWH whose NK cells were able to either significantly reduce p24 levels compared to baseline (good responders) or remained dysfunctional an unable to kill infected cell (bad responders) after stimulation with Nano-PIC DCs (Figure 2A). The majority (78.6%) of good responder PWH exhibited a reduction over 50% of p24+ cells after co-culture with NK-NanoPIC-DC. Similarly, 64.3% of individuals from the bad responder PWH group were characterized by exhibiting over 50% increase in p24+ cells after NK-NanoPIC-DC stimulation (Supplemental Table 3). We then analyzed potential differences in clinical data from good and bad responders (Supplemental Table 1). No differences were observed in terms of gender (100 and 91% males), time since ART initiation (7 years versus 13 years), NADIR CD4+ T cell counts (452 and 362), CD4+ vs CD8+ T cell ratios (1.07 and 0.89) and plasma viral load at diagnosis (when available, 96,000 (n=8) and 53,000 (n=9) between good and bad responder PWH groups, respectively. However, we observed a significant difference of lower age (p=0.01) and a trend towards higher CMV coinfection (78% versus 62%; p=0.07) in the good responder PWH group (Supplemental Table 1). In addition, no significant differences were observed in the proportions of total CD56dim and CD56lo/- CD16+ NK cell subsets after Nano-PIC DC between the two PWH groups (Supplemental Figure 3C).

**Figure 2.**
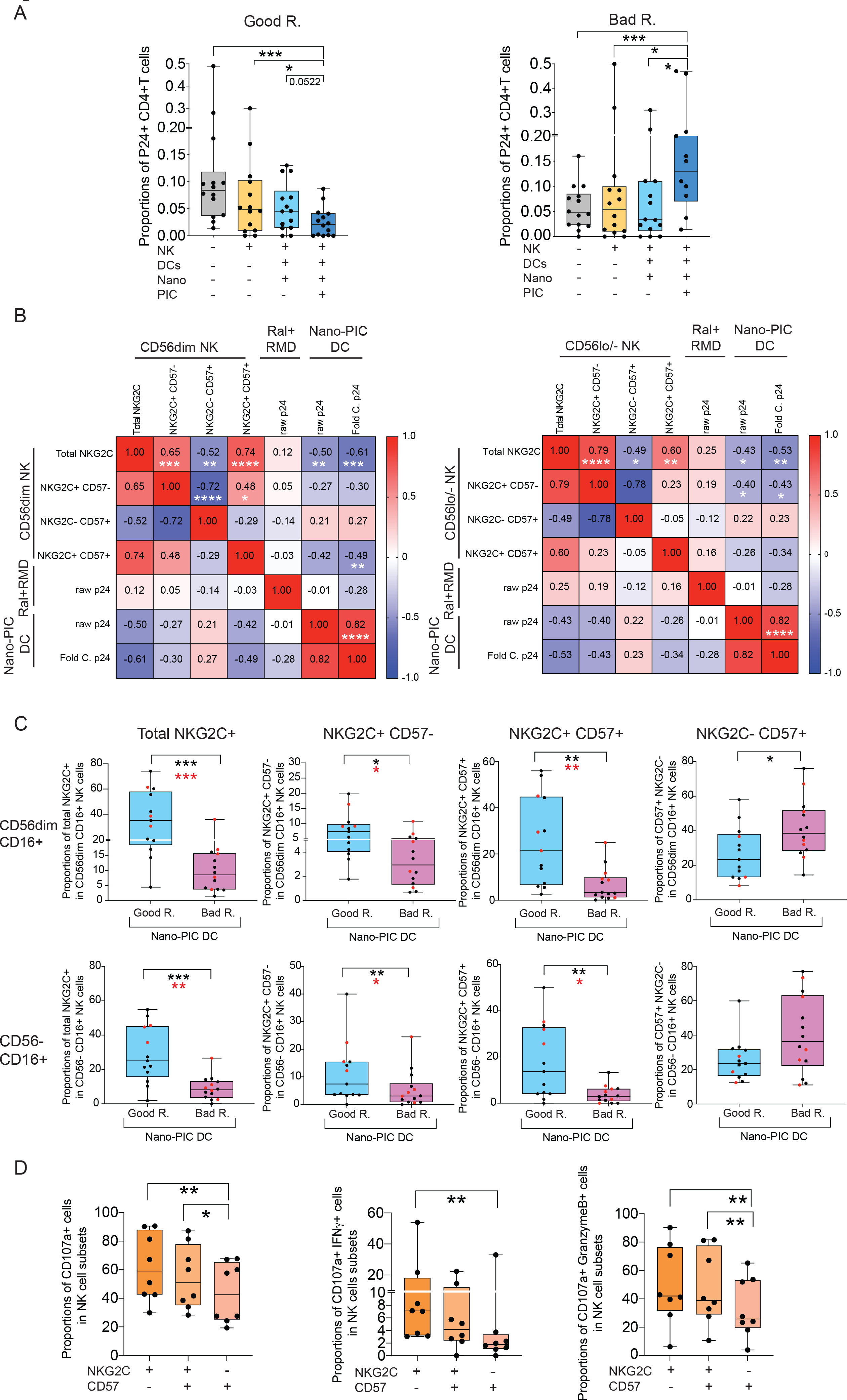
Impact of Nano-PIC DCs on cytotoxic function of NK cells from PWH against HIV- 1 infected CD4+ T cells. (A): Analysis of intracellular expression of HIV-1 p24 on CD4+ T cells from aviremic ART-treated PWH treated with Raltegravir and Romidepsin, in the absence or the presence of NK cells alone or stimulated with DCs treated with empty Nano or Nano-PIC. Data from two separate groups with good (Good R.) and bad (Bad R.) functional response to Nano-PIC DC are shown in the left (n=14) and right (n=14), respectively. (B): Heatmaps representing Spearman correlation matrix between the indicated memory NK subsets within CD56dim (left) and CD56lo/- (right) and proportions of p24+ cells at baseline or after coculture with NK- NanoPIC-DC immunotherapy. Fold changes in p24+ frequencies are also included. Levels of positive and negative associations are highlighted in different intensities of red and blue, respectively. Significant associations have also been highlighted. *p<0.05; **p<0.01; ***p<0.001; ***p<0.001. (C): Proportions of total NKG2C+ cells and different NK subsets based on combination of NKG2C and CD57 on CD56dim CD16+ (upper plots) and CD56lo/- CD16+ (lower plots) NK subpopulations from good (n=13, Good R.; blue) and bad (n=13, Bad R; pink) responder PWH. Good and bad responder PWH not reaching a fold change in p24+ frequencies <0.5 or >1.5 have been highlighted in red. (D): Proportions of total CD107a+ (left), IFNγ+ CD107a+ (middle) and CD107a+ Granzyme B+ (right) cells included in gated memory NK precursors NKG2C+ CD57-, mature memory NKG2C+ CD57+ and effector NKG2C- CD57+ NK subsets from (n=8) PWH after PMA and Ionomycin stimulation (see methods). Statistical significance was calculated using a two tailed-Wilcoxon test. P values calculated excluding individuals not reaching mentioned criteria are also shown in red. *p<0.05; **p<0.01; ***p<0.001.

We then assessed whether reinvigoration of cytotoxic NK cells from PWH by NanoPIC-DC defined as decrease in proportions of p24+ cells may be associated with the induction of adaptive NKG2C+ NK cells, which have been linked to the control of HIV-1 infection (*70*). Moreover, we also analyzed co-expression of NKG2C with CD57, which allows to distinguish between fully mature NKG2C+CD57+ memory and NKG2C+ CD57- precursor memory NK cells (*71*). We observed a significant negative correlation between proportions of total NKG2C+ and NKG2C+ CD57+ NK within CD56dim CD16+ NK cells and percentages of p24+ cells (Figure 2B). Consistently, significant higher levels of total NKG2C+ cells and mature NKG2C+ CD57+ NK cells responders were found within CD56dim CD16+ and also in CD56negative CD16+ cell subsets from PWH good compared to bad responder PWH (Figure 2C). Interestingly, we also observed that NKG2C+ CD57- memory precursor cells were also increased both in CD56dim and CD56lo/- CD16+ NK subsets in good responder PWH compared to bad responders (Figure 2C). In contrast, effector NKG2C- CD57+ NK cells were significantly and negatively correlated with total NKG2C+ cells and were enriched within CD56dim and CD56lo/- NK subpopulations in bad compared to good PWH responder groups (Figure 2B-C). Importantly, these differences essentially remained significant when considering the most extreme good and bad responder phenotype (at least 50% reduction or 50% increase in p24+ percentages, respectively (Supplemental Table 3) (Figure 2C, excluding red dots). Interestingly, similar tendencies in memory and effector NK subsets also appeared to be present in the two PWH groups at baseline in the absence of DC stimulation (Supplemental Figure 3D). Finally, NKG2C+ CD57- memory NK cell precursors were characterized by higher expression of degranulation markers such as CD107a, and higher proportions of CD107a+ cells co-expressing IFNγ were found compared to effector NKG2C- CD57+ NK cells (Figure 2D). Likewise, higher percentages of total IFNγ+ cells as well as increased CD107a+ granzyme B+ were also found in NKG2C+ CD57- memory precursor cells (Figure 2D, Supplemental Figure 3E). In contrast, levels of TNFα tended to be lower on NKG2C+ CD57- NK cells and higher on NKG2C- CD57+ effector NK and total granzyme B expression was almost 100% in all populations (Supplementary figure 3E). Together, these findings suggest that effective response of NK cells to Nano-PIC DC immunotherapy is associated with the induction of memory NKG2C+ NK cells, which may have improved capabilities to target HIV-infected CD4+ T cells.

### Cytotoxic function of NK cells from PWH primed with Nano DC-PIC is mediated through TRAIL

We next investigated the mechanism of enhanced cytotoxicity observed in NK primed with Nano- PIC DCs. To this end, we analyzed the expression of different ligands for NKG2C, NKG2D and TRAIL receptors on HIV-1 infected CD4+ T cells from PWH (Supplemental figure 4A). As shown in Figure 3A, higher levels of the NKG2D ligand MICa/b, the NKG2C ligand HLA-E, and the TRAIL ligand DR4 were enriched on reactivated p24+ CD4+ T cells from PWH. Interestingly, CD56dim CD16+ and CD56lo/- CD16+ NK cells from good responder PWH tended to express higher levels of TRAIL compared to bad responder individuals at baseline and after Nano-PIC DC stimulation (Supplemental Figure 4B-C). In this regard, these differences seemed to be more marked in both memory NKG2C+ CD57- precursor population which expressed significantly higher levels of TRAIL compared effector NKG2C- CD57+ cells in both CD56dim and CD56neg CD16+ subsets from good responder PWH (Figure 3B). Expression of TRAIL also tended to be higher in mature NKG2C+ CD57+ memory NK included in CD56lo/- CD16+ cells from good versus bad responder PWH (Supplemental figure 4D). To determine whether killing of NK cells primed with Nano-PIC DC was dependent on TRAIL or other potential activating receptors such as NKG2C and NKG2D, we performed functional co-culture assays of NK primed with Nano-PIC DC and autologous CD4+ T cells from good responder PWH in the presence of blocking mAbs directed to each of these molecules or control mAb. Notably, blockade of TRAIL selectively impaired the reduction of p24+ CD4+ T cells by autologous NK and Nano-PIC DC (Figure 3C). In contrast, a partial non-significant inhibition of the reduction of p24+ cells was observed after the blockade of NKG2C or NKG2D, suggesting these receptors were not involved in the main killing mechanism participating in our model (Figure 3C). Together, these data indicate that enhanced functionality of NK cell and improved elimination of HIV-1 infected cells after DC-based immunotherapy is associated with TRAIL expression, which is enriched on NK cells from good responder PWH and enriched in NKG2C+ memory NK cells.

**Figure 3.**
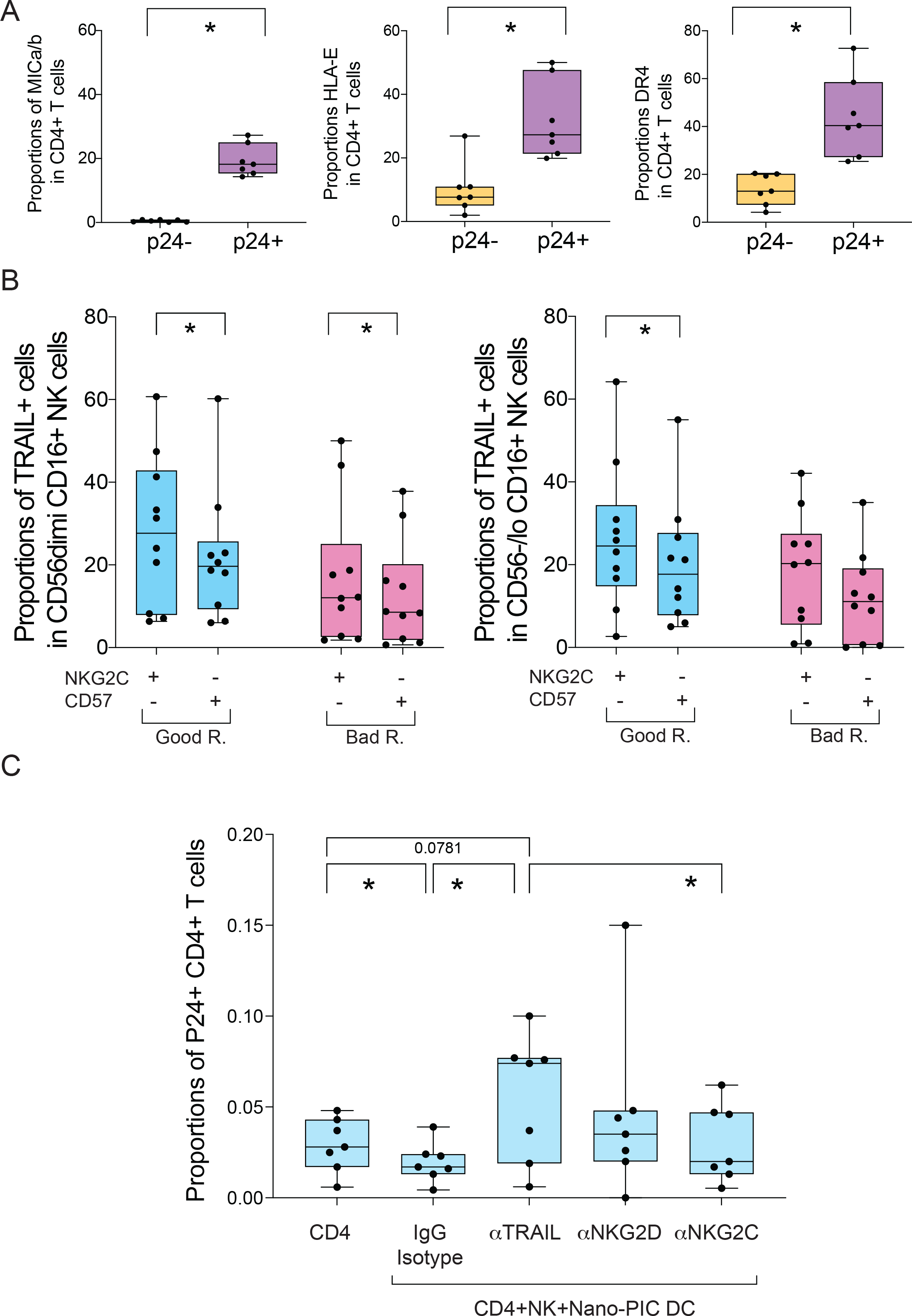
Analysis of expression of NK receptor-ligands and mechanisms of cytotoxic function of NK cells primed with Nano-PIC DC. (A): Proportions of CD4+ T cells from PWH expressing the NK receptor ligands MICa/b (left), HLA-E (middle) and DR4 (right) within HIV- 1 p24+ (purple) and p24- (yellow) populations in n=7 PWH. (B): Analysis of proportions of cells expressing TRAIL in NKG2C+ CD57- and NKG2C- CD57+ cell subsets from good (n=10, Good R.; blue) and bad (n=10, Bad R; pink) responder PWH included in CD56dim CD16+ and cd56lo/- CD16+ NK subpopulations. (C): Proportions of p24+ CD4+ T cells from n=7 PWH previously identified as good responders treated with Raltegravir and Romidepsin and cultured in the absence or the presence of autologous NK cells treated with Nano-PIC DCs and either IgG Isotypic control or with anti-NKG2D, anti-NKG2C or anti-TRAIL blocking antibodies. Statistical Significance between experimental conditions in the same samples or between different study cohorts were calculated using a two tailed Wilcoxon matched pairs or a Mann Whitney tests, respectively. *p<0.05.

### Expression of TIGIT limits induction of functional memory NKG2C+CD57- NK and cytotoxic function against HIV-infected cells

To determine whether different levels of distinct checkpoint receptors could also be associated with the differential enrichment on memory and effector CD56dim CD16+ and CD56lo CD16+ NK observed in good and bad responder PWH we analyzed the expression of PD-1, TIGIT and TIM3 in these populations after stimulation with Nano-PIC DC (Supplemental figure 5A). No associations between PD-1 expression on CD56dim and CD56lo/- CD16+ NK populations were found in good and bad responder PWH after treatment with Nano-PIC DC (Supplemental Figure 5B-C). In contrast, TIGIT and TIM3 appeared to be differentially and inversely enriched in CD56dim and CD56lo/- NK populations from the two PWLH subgroups (Supplemental Figure 5B-C). In fact, TIGIT expression was associated with lower percentages of NKG2C+ memory NK cells within the CD56dim CD16+ population and with higher frequencies of effector CD57+ NKG2C- cells in CD56lo/- CD16+ NK cells (Figure 4A). In contrast, TIM3 expression was significantly correlated with the frequencies of NKG2C+ CD57- memory precursor NK cells in both CD56dim CD16+ and CD56lo/- CD16+ populations (Figure 4A). Interestingly, TIM3 expression also showed a highly significant correlation with TRAIL expression on both CD56dim and CD56lo/- NK subpopulations (Figure 4A). Moreover, we also found that expression of TIGIT was also higher in the few NKG2C+ CD57- precursors present in CD56dim and CD56lo/- NK subpopulations from bad responder PWH (Figure 4B). Similar trends were observed for the more mature memory NKG2C+ CD57+ and effector NKG2C- CD57+ cells (Figure 4B, Supplemental Figure 5D). Therefore, TIGIT and TIM3 differentially associate with the induction of functional versus exhausted NK cell subsets in response to Nano-PIC DC immunotherapy.

**Figure 4.**
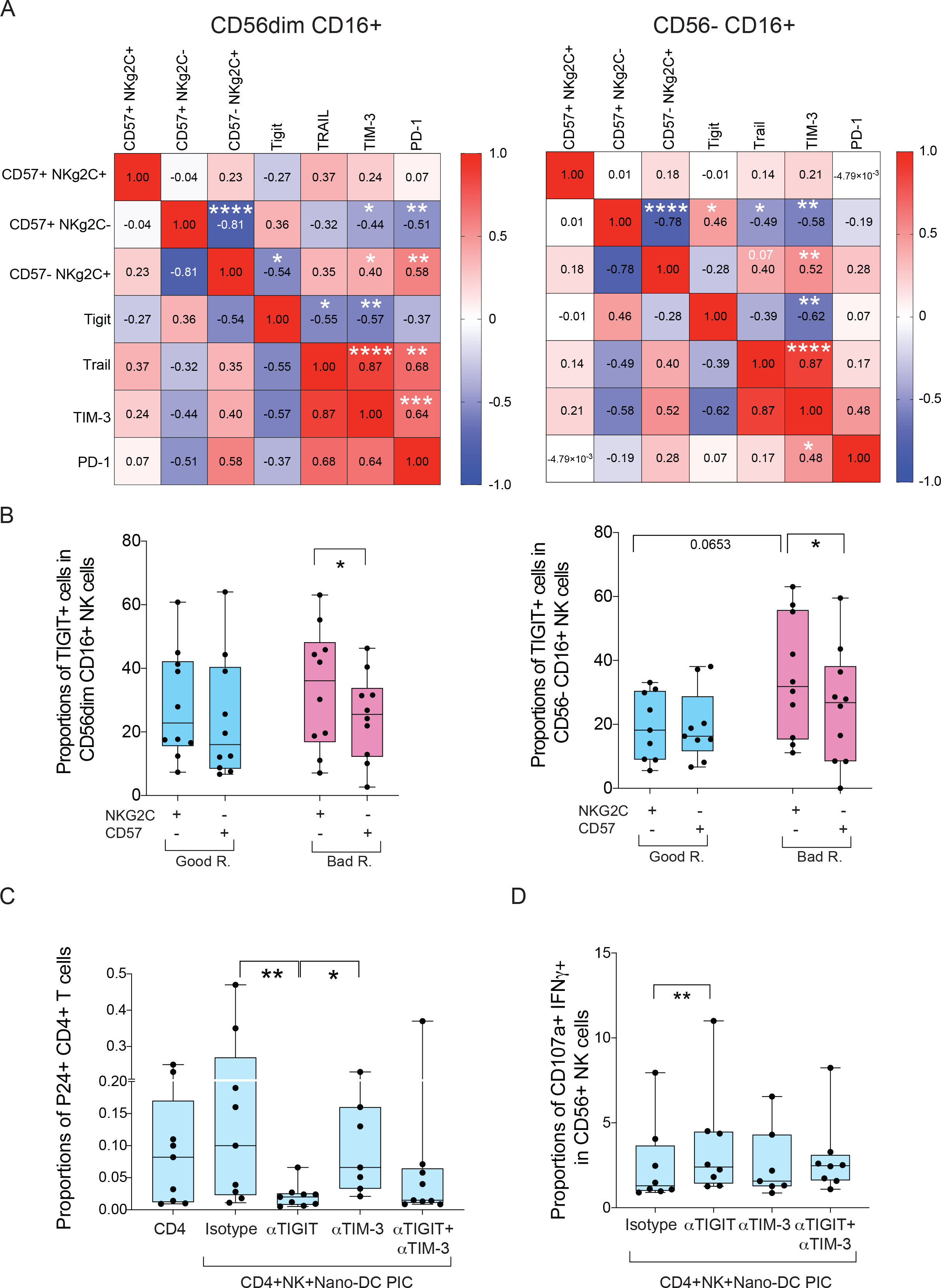
Association of expression checkpoint receptors with proportions and phenotype of memory and effector NK cell subsets from PWH in response to Nano-PIC DC. (A): Heatmaps representing spearman correlation networks between proportions of NKG2C+ CD57-, NKG2C+ CD57+ and NKG2C- CD57- cell subsets, expression of TRAIL and checkpoint receptors TIGIT, PD-1 and TIM3 within CD56dim CD16+ and CD56lo/- CD16+ NK populations from all PWH included in the study. (B): Proportions of TIGIT+ cells within memory NKG2C+ CD57- and effector NKG2C- CD57- cell subsets from good (n=10; Good R.; blue) and bad (n=10; Bad R; pink) responder PWH included in CD56dim CD16+ and cd56lo/- CD16+ NK subpopulations. (C): Proportions of CD4+ T cells expressing HIV p24 cultured alone or with NK treated with Nano- PIC from n=9 bad responder PWH in the presence of isotypic control mAb or individual or combined treatment with anti-TIGIT. Anti-TIM3 mAbs were also included in n=7 experiments. (D): Proportions of CD107a+ IFNγ+ NK cells in the co-culture experimental conditions described in (C). Statistical significance was calculated using a two tailed-wilcoxon test (B,C,D) and statistically significant Spearman associations are highlighted in the heatmap (A). *p<0.05; **p<0.01; ***p<0.001; ****p<0.0001.

Next, we determined whether blocking TIGIT could restore the cytotoxic properties of NK cells that remained dysfunctional in response to Nano-PIC DC from the bad responder PWH group. To this end, we performed additional functional assays using NK cells from this PWH group stimulated with Nano-PIC DC and analyzed their ability to eliminate autologous reactivated p24+ CD4+ T cells in the presence of anti-TIGIT or anti-TIM3, or the combination of both blocking Abs or the corresponding Isotypic control Abs. As shown in Figure 4C, blockade of TIGIT led to a marked reduction in the frequencies of p24+ CD4+ T cells in the presence of NK cells primed with Nano-PIC DC suggesting the restoration of NK cell function. In contrast, treatment with isotypic control Abs or TIM3 antibodies did not affect frequencies of infected cells, suggesting the effect was specific of TIGIT. Supporting an increase of cytotoxic function of NK cells in the presence of anti-TIGIT mAbs, a higher proportion of cells co-expressing CD107a and IFNγ were found in these conditions but not in the presence of isotypic control Abs, anti-TIM3 or the combination of both anti-TIGIT and anti-TIM3 antibodies (Figure 4D). In addition, we discarded that the anti-TIGIT mAbs had an impact on frequencies of p24+ cells when added to cultures of isolated CD4+ T cells from PWH cultured individually in the presence of media or after reactivating with PHA, suggesting that they did not significantly revert viral latency or mediate elimination of infected cells (Supplemental Figure 5E). However, we did observe expression of the ligand of TIGIT in reactivated p24+ CD4+ T cells from some PWH tested, supporting that blockade of this interaction may also have an impact in the elimination of these cells (Supplemental Figure 5F). Thus, our data demonstrate that blockade of TIGIT can improve functional restoration of NK cells from bad responder PWH in combination with Nano-PIC DCs.

### Combination of Nano PIC-DC/NK immunotherapy and blockade of TIGIT reduces expansion of HIV-1 infected cells and preserves TRAIL+ Granzyme B+ NKG2C+ memory NK in vivo

To better understand the relevance of NKG2C+ memory NK cell precursors to control HIV-1 infection, we first analyzed the dynamics of these cells *in vivo* in a humanized BLT mouse model, which allows the reconstitution of human immune cells including NK cells and their maturation into CD16+ cells upon injection with rhIL-15 (Supplemental Figure 6A). No significant changes in body weight were observed during the course of experiment in these hBLT mice (Supplemental Figure 6B). When we analyzed the reconstitution of these animals (Supplemental Figure 6C), a significant expansion of human cells and mature CD16+ CD56+ NK cells was observed upon injection with rh IL-15 (Supplemental Figure 6D-E). We then compared evolution of NK cells in hBLT mice infected with HIV-1 compared with an uninfected group of animals. As expected, infected hBLT displayed detectable plasma HIV-1 viral loads peaking at 2 weeks post-infection and a subsequent reduction in proportions of CD4+ T cells within total CD3+ cells at 3 weeks (Supplemental figure 6F) in agreement with kinetics reported in previous studies with this model (*69*). An early expansion of NKG2C+ CD57- memory NK precursors within CD56dim CD16+ population tended to occur at 1 week post-infection in infected mice, and this subset was subsequently reduced at 2 and 3 weeks and progressively enriched in the CD56lo/- NK population at these times post-infection (Supplemental figure 6G). Interestingly, proportions of fully mature NKG2C+ CD57+ NK within both circulating CD56dim and CD56lo/- CD16+ NK cells followed opposite dynamics and were progressively increased after HIV-1 infection specially at 2 weeks and subsequently disappeared from the blood by 3 weeks (Figure 5A-B). Importantly, fully mature NKG2C+ CD57+ NK cells associated with higher plasma viremia at weeks 1, 2 and 3 post infection (Figure 5C, left heatmap). In contrast, preserved frequencies of NKG2C+CD57- memory precursor NK cells significantly correlated with less severe depletion of CD4+ T cells in PB from transplanted NSG mice (Figure 5C, right heatmap). In addition, effector NKG2C- CD57+ NK cells accumulated in CD56lo/- CD16+ population from HIV-1 infected hBLT mice (Supplemental figure 6G). Thus, as we previously observed *in vitro* assays, preserved memory NKG2C+ cell precursors may associate with preserved NK function *in vivo*.

**Figure 5.**
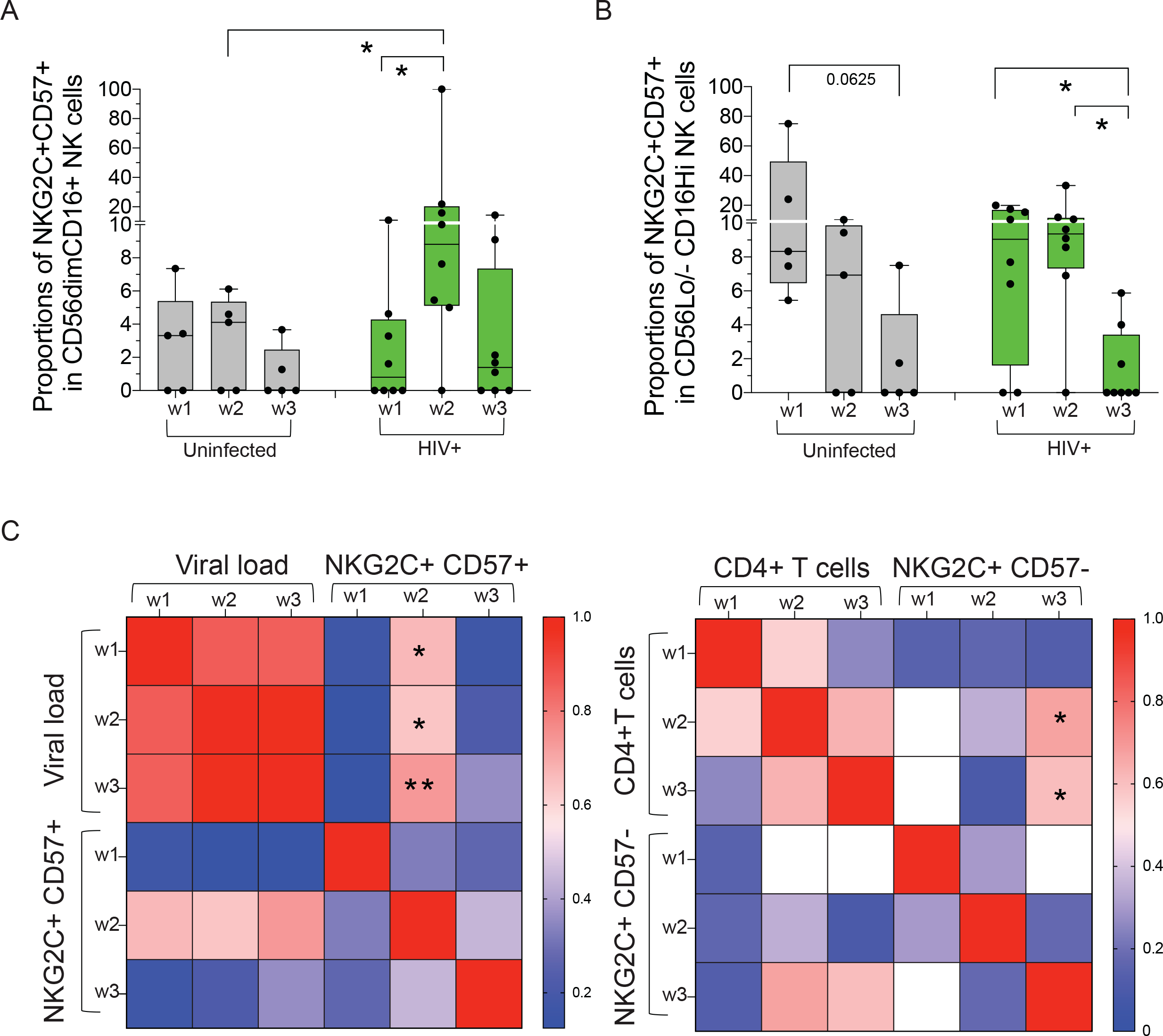
Mature memory NKG2C+ CD57+ NK cells associate with viral load in humanized BLT mice during HIV-1 infection. (A,B): Proportions of mature memory NKG2C+ CD57+ within circulating CD56dim CD16+ (A) and CD56lo/- CD16+ (B) NK populations in n=8 hBLT mice after 1, 2 and 3 weeks of infection with HIV-1 JRCSF (green). Data from n=5 uninfected hBLT mice (grey) analyzed in parallel at the same time points are also shown. Statistical significance was calculated using a two tailed-wilcoxon test. *p<0.05 (C): Spearman correlation networks showing correlations between proportions of mature memory (left) and memory precursors (right) NK cells and effector NK cells with plasma HIV-1 viral load and CD4+ T cell proportions in HIV-1 infected hBLT mice, respectively. Positive and negative correlations and intensity are highlighted in red and blue, respectively. Statistically significant spearman associations are highlighted in the heatmap. *p<0.05; **p<0.01.

Finally, we aimed to assess whether TIGIT blockade could potentiate Nano-DC PIC and NK cell immunotherapy and elimination of HIV-1 infected CD4+ T cells *in vivo* in the absence of CD8+ T cells. To this end, we used a different humanized mouse model based on immunodeficient NSG mice transplanted with pre-isolated CD4+ T cells from PWH by intravenous infection. Three independent *in vivo* experiments using cells from three different PWH donors were performed (Supplemental Table 4). Two days later, mice were split in three different groups of either animals left untreated to allow expansion of human infected CD4+ T cells or injected intravenously with autologous Nano-DC PIC and NKs and receiving intra-peritoneal injections of either anti-TIGIT or Isotype mAbs (Figure 6A). The blocking anti-TIGIT (Tiragolumab) and isotype control antibodies were then administered every two days for 7 days (Figure 6A). Detection of viremia in the NSG mouse viral outgrowth model (MVOA) is quite variable and may rely on the initial frequencies of infected CD4+ T cells transplanted, the level of expansion of these cells mediated by mechanisms such as GvHD. In agreement with these previous studies (*72*), human CD45+ cells from PWH were detected in the blood and in the spleen of xenotransplanted NSG mice (Supplemental Figure 7A-B). Interestingly, a preferential distribution of human cells in the spleen versus the PB was observed in mice treated with anti-TIGIT mAbs in contrast to those receiving isotype controls (Supplemental Figure 7B). Consistently, numbers of hCD4+ T cells were also significantly higher in the spleen in animals treated with Nano-PIC DC and NK cells and anti-TIGIT mAbs (Supplemental figure 7B). As expected, we were not able to detect HIV-1 viral loads over the limit of detection in the plasma in transplanted NSG mice characterized by low reconstitution of human cells (Supplemental figure 7C). Alternatively, to assess potential effects on the selective elimination of HIV-1 infected CD4+ T cells in different body compartments, we analyzed the intracellular expression of p24 in in human CD4+ T cells form peripheral blood and spleen cells in infected NSG mice by flow cytometry (Supplemental Figure 7D). As shown in Figure 6B, we detected a considerable increase of up to a median of 23.35 % of p24+ cells in human circulating CD4+ T cells from the transplanted NSG mice untreated with NK-NanoPIC DC immunotherapy, suggesting an expansion of infected cells in periphery. Moreover, the treatment of mice with Nano-PIC DC and NK cells significantly reduced proportions of p24+ cells within circulating CD4+ T cells, suggesting that immunotherapy by itself was able to control/reduce expansion of infected cells in PB (Figure 6B). Although no significant differences were found in median levels of p24 in the PB between mice groups treated with anti-TIGIT and isotypic control Ab, a higher proportion of animals with undetectable p24 expression within CD4+ T cells was present in the anti-TIGIT group (Figure 6B, left). When we analyzed levels of p24+ cells in tissue, we again observed a higher number of mice that displayed undetectable levels of infection in the NSG treated with NK-NanoPIC-DC immunotherapy and the anti-TIGIT Antibody (Figure 6B, right). The MVOA model allows the visualization of cluster on human HIV-1-infected cells in the lymphoid tissue of transplanted mice (*72*). To corroborate whether differences in frequencies of p24+ cells in mice treated with anti-TIGIT mAbs could be associated to control of spread of infected cells, we assessed the size of p24+ cell clusters in the spleen by histology of the different transplanted NSG mice groups (Figure 6C, left; Supplemental figure 7E). Notably, we observed a significant reduction in the size of clusters in the NSG mice group treated with Nano-PIC DC-NK immunotherapy and the anti-TIGIT mAb (Figure 6C, right).

**Figure 6.**
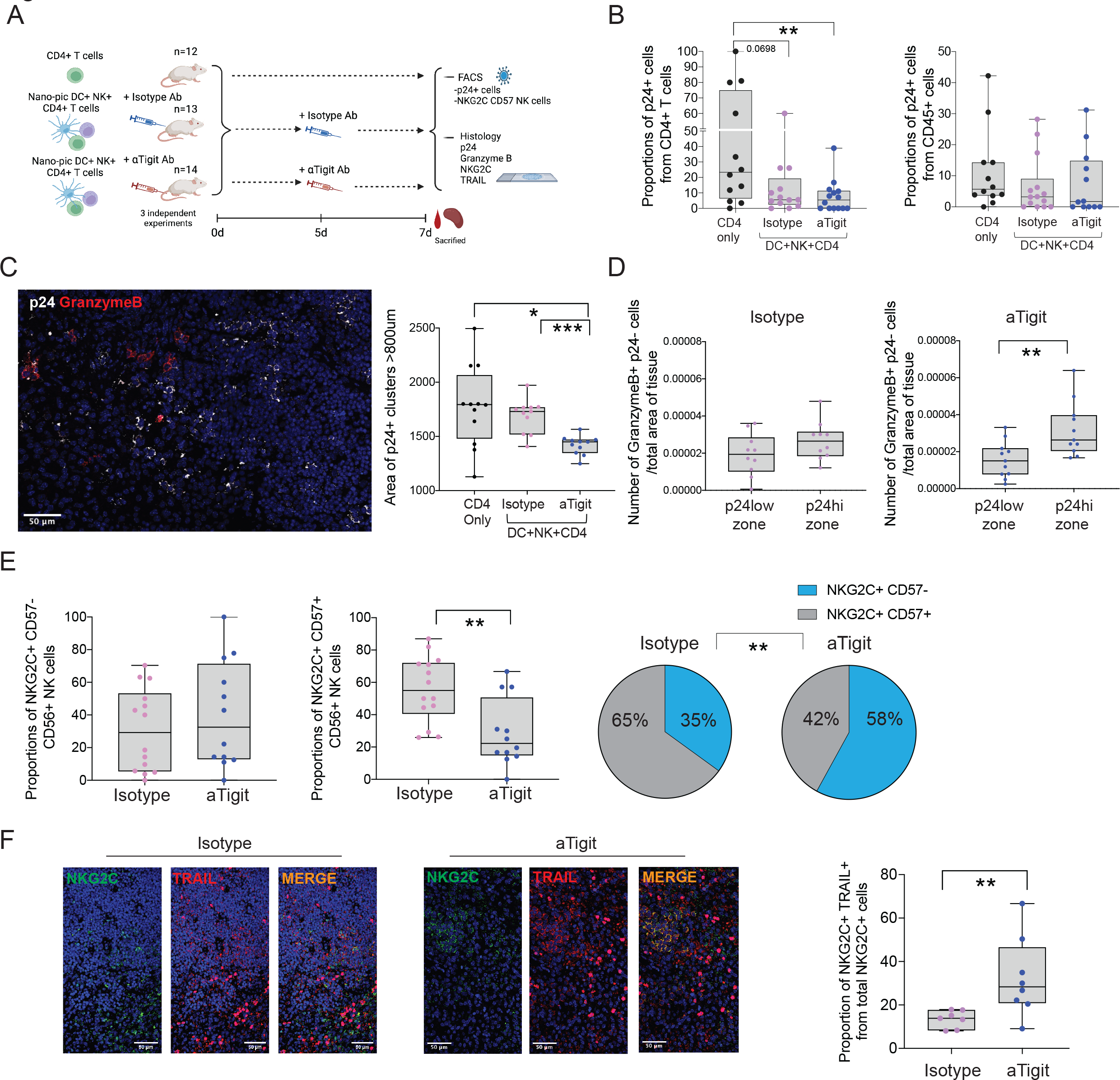
Blockade of TIGIT enhances NK-NanoPIC-DC immunotherapy against HIV-1- infected CD4+ T cells from PWH *in vivo* in transplanted NSG mice (A): Schematic representation of experimental humanized NSG mouse model and the different NK-NanoPIC-DC immunotherapy, and treatment with either Isotypic control and anti-TGITmAb. Number of total combined mice from n=3 independent experiments. (B): Proportions of HIV-1 p24+ cells within CD4+ T cells in the peripheral blood (n=12, left plot) and spleen (n=12, right) of NSG mice transplanted with CD4+ T cells from PWH alone (black) or in combination with autologous NK and Nano-PIC DC and injected with either Isotypic (n=13 blood and n=14 spleen; violet) or anti-TIGIT (n=14 blood and n=12 spleen; Blue,). (C): Analysis of size of clusters of p24+ cells present in the spleen of transplanted NSG mice with CD4+ T cells from PWH alone (n=12; black) or in combination with autologous NK and Nano-PIC DC and injected with either isotypic (n=11; violet) or anti-TIGIT (n=11; blue). A representative confocal microscopy image showing expression of p24 (white) and granzyme B (red) is shown in the left. (D): Quantification of granzyme B+ cells around and within areas of low and high p24+ clusters in spleen from transplanted NSG mice receiving NK-Nano-PIC and isotype (n=10; violet) or anti-TIGIT (n=11; blue) immunotherapy corrected by the total area of tissue analyzed. (E): Proportions of NKG2C+ CD57- precursors (violet) and NKG2C+ CD57+ mature memory NK cells from hCD45+ CD56+ cells present in the spleen of humanized NSG mice treated with isotype (n=14; violet) or anti-TIGIT (n=12; blue) antibodies. Quantification of proportions of each population within the total NKG2C+ population is also shown on the right. (F): Quantification NKG2C and TRAIL colocalization in the spleen of humanized NSG mice treated with isotype (n=7; violet) or anti-TIGIT (n=8; blue) antibodies is shown in the left. Representative 40X magnification confocal microscopy images showing co-expression of NKG2C (green) and TRAIL (red) individually and merged in the spleen of NSG mice receiving anti-TIGIT and isotypic control Abs. Statistical significance between different experimental groups was calculated using either a two tailed Mann whitney test or a Chi-square test. *p<0.05; **p<0.01; ***p<0.001.

Finally, we analyzed whether there was an association between the maturation and cytotoxicity of memory NK cells and the reduction of p24+ cells in the spleen of transplanted NSG treated with anti-TIGIT mAbs. We observed significantly higher numbers of granzyme B+ p24- cells around areas of high p24 expression in the spleen of mice treated with anti-TIGIT antibodies (Figure 6D, Supplemental Figure 7E). We then analyzed proportions of NKG2C+ memory NK in these tissues (Supplemental Figure 7F). Interestingly, we also detected a significant expansion in frequencies of total memory NKG2C+ cells in two out of three experiments in transplanted mice as compared to baseline levels present prior to injection (Supplemental Figure 7G). We observed that fully mature memory NKG2C+ CD57+ NK cells were enriched in the animals treated with immunotherapy and Isotype mAb control compared to the group treated with the anti-TIGIT mAb, which resulted in a significant enrichment of NKG2C+ CD57- precursors within the total pool of memory NK cells in the spleen after anti-TIGIT mAb treatment (Figure 6E; Supplemental Figure 7F). Last, a higher proportion of NKG2C+ cells detected in the spleen from these mice co-expressed TRAIL (Figure 6F). Together, these results indicate that the combination of Nano-DC PIC and NK cells immunotherapy with blockade of TIGIT more efficiently reduces the spread of HIV-1 CD4+ T cells *in vivo*, and associates with preserved frequencies of NKG2C+ CD57- precursors, increased cytotoxic profiles characterized by higher expression of Granzyme B and TRAIL in the tissue.

## DISCUSSION

The achievement of a remission of HIV-1 infection requires either the complete elimination or significant reduction of persistent HIV-1 reservoirs leading to either a sterilizing or functional cure of the infection, respectively. To this end, multiple strategies are being explored including trapping remaining viral reservoirs and preventing new replication (block and lock) (*73, 74*) or to potentiate the immune response against latently infected cells (Shock and Kill, Kick and kill) ((*75–77*). Previous studies have shown that modulation of the immune responses against HIV-1 in PWH even on ART has in many cases not been successful due to an exhausted hyporesponsive state of immune cells in PWH. This is characterized by increased expression of checkpoint receptors and the lack of response upon antigenic or cell to cell stimulation (*78–80*). The present study addresses the potential of DC immunotherapies increasing cytotoxic activity of NK cells from PWH against autologous HIV-1 infected CD4+ T cells. Our data show that modulation of DC with nano-PIC increased their ability to promote the expression of cytotoxic markers and natural cytotoxicity function on NK cells *in vitro*, and that the signals provided by DC to NK cells require cell-to-cell contact. Hence, DC treated with Nano-PIC induce expression of the NKG2D ligands MICa/b and UBLP1, thereby supporting a potential interaction mediated via NKG2D-ligand. In a different physiopathological scenario, previous data showed that activation of the intracellular sensor RIG-I induces expression of the NKG2D ligand MICa/b on cDC2 and this interaction is needed to induce aberrant activation of NK cells in the autoimmune Sjogreńs Syndrome (*68*). In addition, although cytokines secreted by DC alone were not sufficient to induce activation of NK cells, it is known that they play a role in NK cell maturation and education (*81, 82*). In fact, IL-15 secreted by DC has been involved in the activation of NK cells (*83*), but in our model the expression of this cytokine was not detected. Instead, Nano-PIC DC secreted higher levels of IL-12 and IFNβ which also contribute to modulate NK cell survival and activation (*84*).

Our data also provide novel information about enhancement of NKG2C memory NK cells in response to DC immunotherapy. While some studies support a protective role of these cells during HIV-1 infection in HIV post-treatment controllers (*62*) and in the SIV-primate model (*85*), there was no previous information regarding the modulation of this NK cell subset in response to DC immunotherapy. In this regard, our data provide new evidence regarding the association of NKG2C+ memory NK cells and the effective elimination of HIV-1 infected CD4+ T cells in functional assays with these cells. Previous reports associated increased proportions of NKG2C+ memory NK cells to CMV infection (*86*). Consistently, we observed a tendency to higher coinfection with CMV in PWH capable of more efficiently inducing memory NKG2C+ cells and improving functionality after exposure to Nano-PIC DC. These findings are consistent with previous studies that reported an expansion of memory NKG2C+ NK during acute CMV and HIV infections that is mainly driven by viremia (*87, 88*). Therefore, CMV co-infection could represent a biomarker of individuals that received a priming on NK cells that could be used as a criterion to select PWH candidates for NK immunotherapy against HIV-1. However, this possibility should be investigated in more detail in preclinical studies.

Several studies have already described that NKG2C+ memory NK cells display higher ADCC function (*89*). Although our results also support increased ADCC activity in the presence of bNAbs, our study identifies a TRAIL-dependent mechanism of natural cytotoxicity potentially also operating in these NKG2C+ memory NK cells. Interestingly, TRAIL has previously been involved in the molecular mechanism allowing elimination of HIV-1 infected CD4+ T cells by NK cells *ex vivo* and controlling the reservoir size in humanized mice (*90–92*) and increased expression of this molecule had been reported in IL-15 expanded NK cells from PWH (*93*). Also, TRAIL has been proposed as a mechanism of killing utilized by tissue-resident NK cells in salivary glands in autoimmune Sjogreńs syndrome (*94*) and autoimmune diabetes (*95*). However, the role of TRAIL had not been previously investigated in NKG2C+ memory NK cells or in the context of immunotherapy against HIV-1.

Another important aspect of this study is the identification of TIGIT and TIM3 as two different checkpoint inhibitory receptors associated with differential functional response of NK from PWH to Nano-PIC DCs. These results agree with previous studies reporting accumulation of dysfunctional TIGIT+ NK cells during chronic HIV-1 infection (*96*). Moreover, TIGIT expression is accumulated in mature memory NKG2C+ CD57+ NK cells from women with chronic HIV-1 infection (*97*). Our present results show that TIGIT is associated with dysfunctional state and with the induction of NKG2C+ memory NK. In addition, we found that blockade of TIGIT improves response of NK cells to Nano-PIC DC and targeting of HIV-1 infected T cells *in vitro* and *in vivo*. However, the molecular mechanisms associated to the selective expression of TIGIT and TIM3 with the functional state of NK cells, or whether the blockade of TIGIT exclusively affects NKG2C+ memory NK cells were not analyzed in this study and deserve further studies.

Finally, our work provided in vivo data on the NSG-MVOA model demonstrating that combined blockade of Nano-DC and NK immunotherapy is capable to limiting expansion of HIV-1 infected CD4+ T cells. In these assays, although we did observe an expansion of p24+ infected cells within circulating human CD4+ T cells from PWH, we were not able to detect HIV-1 viral loads in transplanted NSG mice, maybe due to the low frequency of human CD45+ cells in the blood. Supporting this possibility, several studies have previously reported that the efficiency of graft of human cells in these animals combined with the via used to infuse the human cells may greatly affect the detection of plasma viral loads (*98–100*). Our study also focuses on investigating the impact of immunotherapy at earlier time points to avoid the impact of development of GvHD mediated by NK cells and T cells from PWH. However, several studies suggest that viral loads become more detectable beyond 1-week post-transplantation (*72*) and therefore, future studies should be conducted to determine whether this parameter could be applicable to our assay. Instead, our study analyzes the clusters of HIV-1 infected cells in lymphoid tissue, which had already been described in the MVOA model (*72*) as a more reliable parameter to determine the level of expansion of clones of infected cells in the spleen of transplanted mice. Using this approach, we have shown that co-administration of anti-TIGIT antibodies and NK-NanoPIC-DC immunotherapy limits the size of clusters of HIV-1 infected cells in the spleen, suggesting that we may be able to limit the spread of virus in this tissue. Additional *in vivo* data of HIV-1 infection using an alternative hBLT humanized mouse model show a correlation of preserved NKG2C+ CD57- memory precursors with reduced plasma viral load and reduced depletion of CD4+ T cells. However, in this model NK cells were not the only immune cell subsets present in this model *in vivo,* and animals were treated with hIL-15 to increase NK cell reconstitution (*101*), which may have also impact the generation of NKG2C+ memory NK cells and also T cells, as suggested in primate models (*102*). Moreover, we have provided new evidence that this phenotype is associated with a higher expression of TRAIL and granzyme B on NKG2C+ cells also present in the spleen of NSG mice. Thus, taking into account these data combined with a higher enrichment in less differentiated NKG2C+ memory precursors, it is tempting to speculate that we were capable of preserving functional memory NK cells in the tissue in response to anti-TIGIT Abs. For instance, it has been described that tissue resident HIV-1 infected CD4+ T cells express high levels of PD- 1 and TIGIT (*103, 104*) and therefore, may be susceptible to elimination or viral reactivation in the presence of tiragolumab. Although we did not observe a consistent trend to viral reactivation of CD4+ T cells from PWH in vitro, future analyses in NSG mice transplanted exclusively with HIV-1 infected CD4+ T cells should be carried out in order to completely discard this possibility. We have also observed that CD155 tends to be more expressed in reactivated p24+ infected CD4+ T cells from PWH, which may represent an additional potential mechanism by which anti-TIGIT may facilitate targeting of HIV-1 infected cells that are more resistant to elimination. However, conflicting results from previous studies (*97, 105, 106*) indicate that the upregulation of CD155 may not be homogeneous. Therefore, whether this is a significant evasion mechanism used by CD4+ T cells should be studied in more detail. In addition, in future clinical trials, the impact of anti-TIGIT antibodies on CD8+ T cells should also be considered. We previously showed that TIGIT also marks dysfunctional CD8+ T cells unable to respond to DC stimulation (*19*) and the fate of these cells in vivo during HIV-1 infection and in response to anti-TIGIT antibodies should be assessed in the future. Interestingly, TIM3 seemed to associate with the generation of NKG2C+ memory NK cells in our study, but the blockade of this receptor did not result in improvement of their cytotoxic function. Therefore, future studies should focus on the impact of TIM3 on NK cell immunotherapy against HIV-1 as this molecule may display different roles in immune cell regulation (*107*). Despite these limitations, our study identified the expression of TIGIT on NK cells in PWH on ART as a new biomarker relevant to identify candidates with potential limited response to DC-based immunotherapy and lower induction of memory NKG2C+ cells and to potentially recommend a blockade of TIGIT in a personalized fashion in this particular group of individuals. Thus, TIGIT may represent an attractive candidate for immune modulation against HIV-1 (*108*). Collectively, our study has identified DC as an useful tool to increase NKG2C+ memory NK cells as cell population with beneficial therapeutic properties and the blockade of selective checkpoint receptors as a modulator of these therapies, and therefore we have expanded the knowledge about the development of new tools to potentiate NK immunotherapy, which will be relevant for the field to develop new strategies to cure HIV-1 infection.

## LIST OF SUPPLEMENTARY MATERIALS

- Supplemental Figure 1
- Supplemental Figure 2
- Supplemental Figure 3
- Supplemental Figure 4
- Supplemental Figure 5
- Supplemental Figure 6
- Supplemental Figure 7
- Supplemental Table 1
- Supplemental Table 2
- Supplemental Table 3
- Supplemental Table 4

## AUTHOR CONTRIBUTIONS

E.M.G., I.S.C, M.C.M., A.A, F.S.M., and M.J.B. developed the research idea and study concept, designed the study and wrote the manuscript.

E.M.G. supervised the study.

I.S.C., M.A.L and M.C.M designed and conducted most experiments. I.S.C, O.P. carried out in vivo experiments.

F.S.M, I.S, L.GF and M.A.M.F provided samples of HIVnegative and PWH donors and access to culture facilities for functional assays, as well as participated in data discussion.

O.P. and C.D.A. provided technical support.

I.S, L.G.F, J.S, C.M.C provided information of clinical parameters during the study.

M.L.T P.F. and J.A provided NSG immunodeficient mice and contributed to in vivo MVOA experiments.

V.B and A.B. provided humanized BLT mice.

C.D.A, M.A.L and I.T. performed phenotypical analysis from in vitro assays.

M.G. contributed to flow cytometry analysis, provided critical feedback and contributed to the preparation of the manuscript

N.S.G. , I.T. and O.P performed viral load quantification in MVOA and hBLT in vivo experiments.

J.G.P and M.L.D. generated and provided HIV-1 virus stocks for in vivo experiments.

M.C.M. contributed to histological analyses.

### ACKNOWLEDGMENTS

E.M.G. was supported by Ramón y Cajal Program (RYC2018-024374-I), the Spanish Agencia Estatal de Investigación RETOS, Generación de conocimiento and consolidation programs (RTI2018-097485-A-I00; PID2021-127899OB-I00; CNS2023-144841), La Caixa Banking Fundation ETI-CureHIV (HR20-00218) and infectious diseases CIBER from ISCIII (CIBERINFECC). C.D.A. was supported by Comunidad de Madrid Talento Program (2017- T1/BMD-5396). M.C.M was supported by La Caixa Fundation ETI-CureHIV (HR20-00218). HR17-00016 grant from “La Caixa Banking Foundation to F.S.M also supported the study. I.S.C was supported by infectious diseases CIBER from ISCIII (CIBERINFECC). J.G.P was supported by La Caixa Health program ETI-CureHIV project (HR20-00218), the CIBERINFECC from the National Health Institute Carlos III, and the project PI22/01120.

