## Supplemental figures and tables for "Combined Dendritic Cell And Anti-TIGIT Immunotherapy Potentiate Trail+ Memory NK Cells Against HIV-1 Infected Cells"

**Supplemental figure 1. Phenotypical characterization and functional NK priming properties of nano-PIC DC.** (A,C,E): Representative flow cytometry gating strategy defining DC and expression of CD86 in combination with intracellular staining of IL-12 and IFNβ (A) or ligands for NK receptors MICab, ULBP1 and HLA-E (C). Gating strategy defining NK cell subsets based on the expression of CD56 and CD16 and a representative dot plot of intracellular expression of IFNγ and CD107a is shown in E. (B): proportions of live viability dye- DCs (left panel) and surface levels of CD86 on DCs (right panel) after 16h of stimulation with either soluble (Sol) or Nanoparticle-encapsulated (Nano) empty or loaded with PIC in n=6 independent experiments with HD. Staining background controls for (A) and (C) are included.

**Supplemental figure 2. Expression of activating and inhibitory receptors on NK cells stimulated with Nano-PIC DC.** (A, B,C): total proportions of cells expressing NKG2C (A; n=6) ), NKG2D (B; n=6) and NKG2A (C, n=4) on CD56dim CD16+ (left panel) and CD56lo/- CD16+ (right panel) NK subpopulations populations cultured individually (grey) or in the presence of DC treated with empty (Nano, blue) or PIC-loaded (red, Nano-PIC) nanoparticles. Statistical significance was calculated using a two tailed Wilcoxon test. *$p<0,05$. (D): Flow cytometry gating strategy used to analyze NK mediated killing of target K562-GFP cell line in the presence of violet cell trace-labelled NK cells. Dead target K562 cells were considered as violet- cells losing GFP and gaining cell death viability dye expression. Statistical significance was calculated using a two tailed Wilcoxon test. *$p<0,05$. (E): Quantification of Antibody-dependent cellular cytotoxicity killing of CHO target cells overexpressing HIV-1 gp120 protein in the presence of a cocktail of HIV-1 specific bNAbs (VCR01; PGT121; 3BNC117) and NK cells individually or stimulated with Nano or Nano-PIC DC. Values were subtracted from baseline levels. Statistical significance was calculated using a two tailed Wilcoxon test. *p<0,05.

**Supplemental figure 3. Functional restoration of NK from PWH eliminating HIV-1 infected CD4+ T cells after treatment with Nano-PIC DC.** (A): Representative flow cytometry dot plots representing intracellular expression of HIV-1 p24 on CD4+ T cells from a PWH at baseline with raltegative and after reactivation with Romidepsin and in the presence of Raltegravir. A staining background control from an HIV negative donor is also shown. (B): Proportions of HIV1 p24+ CD4+ T cells from all N=28 PWH recruited for the study cultured with Romidepsin and Raltegavir in the absence or the presence of autologous NK cells alone or stimulated with Nano- or Nano-PIC DCs. (C): Proportions of CD56dim and CD56lo/- CD16+ subsets from total NK cells at baseline and after stimulation with Nano-PIC DCs in PWH with effective (n=14, Good R; blue) and defective (n=14, Bad R; pink) functional restoration eliminating HIV-1 infected CD4+ T cells. (D): Proportions of NKG2C+ CD57- memory precursors; mature NKG2C+ CD57+ memory NK cells and NKG2C- CD57+ effector cells in Good (Good R.; blue) and bad (Bad R; pink) responder PWH at baseline. (E): Proportions of total FNγ+ (left), Granzyme B+ (middle) and TNFα+ (right) included in gated memory NK precursors NKG2C+ CD57-, memory differentiated NKG2C+ CD57+ and effector NKG2C- CD57+ subsets from n=8 PWH after 4h stimulation with PMA and Ionomycin in the presence of Brefeldin A and Monensin. Statistical Significance between experimental conditions in the same samples or between different study cohorts were calculated using a two tailed Wilcoxon matched pairs or a Mann Whitney tests, respectively. *p<0.05; **p<0.01; ***p<0.001.

.

**Supplemental Figure 4. Analysis of TRAIL on NK cell subsets from PWH with different response to stimulation with Nano-PIC DC**. (A): Representative flow cytometry dot plots for the NK receptor ligands DR4 (TRAIL), MIC/ab (NKG2D) and HLA-E (NKG2C/A) in gated CD4+ T cells from PWH. The same gating strategy was applied to p24+ and p24- populations. (B): Representative Flow cytometry dot plot showing expression of TRAIL vs NKG2C on gated NK cells from a good responder PWH. (C,D): Proportions of total expression of TRAIL (n=10; good R.; n=10; bad R, C) or proportions of TRAIL+ cells in mature NKG2C+ CD57+ memory NK cells (n=10; good R.; n=10; bad R D) within CD56dim CD16+ (left panel) and CD56-/lo CD16+ (right panel) populations from PWH displaying good (Good R.; blue) and bad (Bad R; pink) response to Nano-PIC DC stimulation. Statistically significant differences between the two groups of PWH were calculated using a two tail Mann Whitney test. *p<0.05.

.

**Supplemental Figure 5. Levels of Checkpoint receptors and TRAIL on NK cells from different groups of PWH**. (A): Representative flow cytometry histograms showing expression of TIGIT, PD-1, TIM3 and TRAIL in gated CD56dim CD16+ NK cells from a representative PWH. Background levels of FMO are overlayed in orange (B,C): Proportions of cells expressing TIM3 (left), TIGIT (middle) and PD-1 (right) on CD56dim CD16+ (B) and CD56lo/- CD16+ (C) NK cells from good (Good R.; blue) and bad (Bad R; pink)  responder PWH in the absence and presence of Nano-PIC DC stimulation. (D): Proportions of TIGIT+ cells within the NKG2C+ CD57+ cell subset included in CD56dim CD16+ (left) and CD56lo/- CD16+ (right) NK populations from good (Good R. Blue) and bad (Bad R. Pink) responder PWH after stimulation Nano-PIC-DC. Statistically significant differences between the two groups of PWH was calculated using a two tail Mann Whitney test. (E): Proportions of p24+ cells in CD4+ T cells from PWH treated with isotypic control (blue) or anti-TIGIT (pink) mAbs in the presence of media alone (Basal) or under reactivation conditions with PHA (PHA). (F): representative flow cytometry dot plots showing expression of PVR (CD155) and HIV p24 in two PWH donors and in a control Healthy donor. Statistical Significance between experimental conditions in the same samples were calculated using a two tailed Wilcoxon matched pairs. *$p<0.05$; **$p<0.01$; ***$p<0.001$.

**Supplemental Figure 6. Characterization of memory NK cell subsets in humanized BLT mice during HIV-1 infection.** (A): Representative flow cytometry dot plots showing identification of human CD45+ cells in reconstituted humanized BLT mice and subsequent gating strategy identifying different myeloid, T cell and NK cell populations. An example of expression of CD16, NKG2C and CD57 on gated human NK cells from hBLT mice is also shown. (B): weigh (g) of individual hBLT mice used during the course of the in vivo experiment before and after HIV-1 infection. (C): Flow cytometry gating strategy from peripheral blood of a representative hBLT mouse identifying hCD45+ cells and discriminating between myeloid CD14+ versus CD3+ T cells. CD4+ T cells are identified within CD3+ cells and NK cells are defined by CD56 vs CD16 expression in CD14-CD3- cells. Expression of NKG2C and CD57 on gated NK is also shown (D,E):

Proportions of human CD45+ (D), total and CD16+ CD56dim NK cells within the pool of human lymphocytes (E) in the peripheral blood of BLT mice before (Wk0) and after 1 and 2 weeks of injection with recombinant human IL-15. (F): HIV-1 plasma viral load (copies HIV-1 RNA/mL; left) and proportions of CD4+ T cells within CD3+ T cells (right) in uninfected and HIV-1 infected BLT mice at 1,2,3 weeks post-infection. (G): Proportions of NKG2C+ CD57- memory precursor cells (upper plots) and NKG2C- CD57+ effector cells (lower plots) included in the CD56dim CD16+ (left) and the CD56lo/- CD16+ (right) NK subpopulations at week 1,2 and 3 post-HIV-1 infection (green) and in the uninfected control mouse group (grey). Statistical significant differences between different mouse groups or in the same animals over time were calculated using two-tailed Mann Whitney and Wilcoxon pair matched tests, respectively. *$p<0.05$; **$p<0.01$; ***$p<0.001$.

**Supplemental figure 7. Analysis of impact of anti-TIGIT antibodies on HIV-1 reactivation and in reconstitution and histological patterns in humanized NSG mice transplanted with NK cells.** (A): Representative flow cytometry gating strategy showing identification of human CD45+ cells from PWH transplanted into immunodeficient NSG mice, and the identification of CD4+ CD3+ T cells. (B): Total absolute numbers of human CD45+ (left,) and CD4+ T cells (right) in the peripheral blood and spleen from NSG mice transplanted either with CD4+ T cells from PWH alone (black) or in combination with autologous NK and Nano-PIC DC and injected with either Isotypic (violet) or anti-TIGIT (blue). Data from n=3 independent experiments is shown. (C): Analysis of HIV-1 plasma viral load on plasma of transplanted NSG mice receiving either only CD4+ T cells from PWH or NK-Nano-PIC-DC immunotherapy and either Isotypic or ant-TIGIT mAbs. A positive control from cells from PWH is also shown. (D): Representative examples of levels of intracellular HIV-1 p24 on gated human CD4+ CD3+ in three extreme examples from transplanted NSG mice. (E): Representative confocal microscopy images showing immunofluorescence histological analysis of a spleen tissue section from a representative transplanted NSG mouse showing staining of HIV-1 p24 (white) and granzyme B (red) showing areas with high p24 cluster concentration. Individual cells with p24 (upper right) and granzyme B (lower right) staining masks are shown (F): Flow cytometry identification of human NK cells from CD3- human lymphocytes based on expression of CD56 and CD16 and expression of NKG2C and CD57 on gated NK cell in transplanted NSG mice (G): Proportions of total memory NKG2C+ NK for each PWH donor used in the three independent *in vivo* experiments at pre-mVOA in after mVOA timepoints. Statistically significant differences between the different groups of treatment was calculated using a two tail Mann Whitney test. *$p<0.05$; **$p<0.01$.

### Supplemental Figure 1

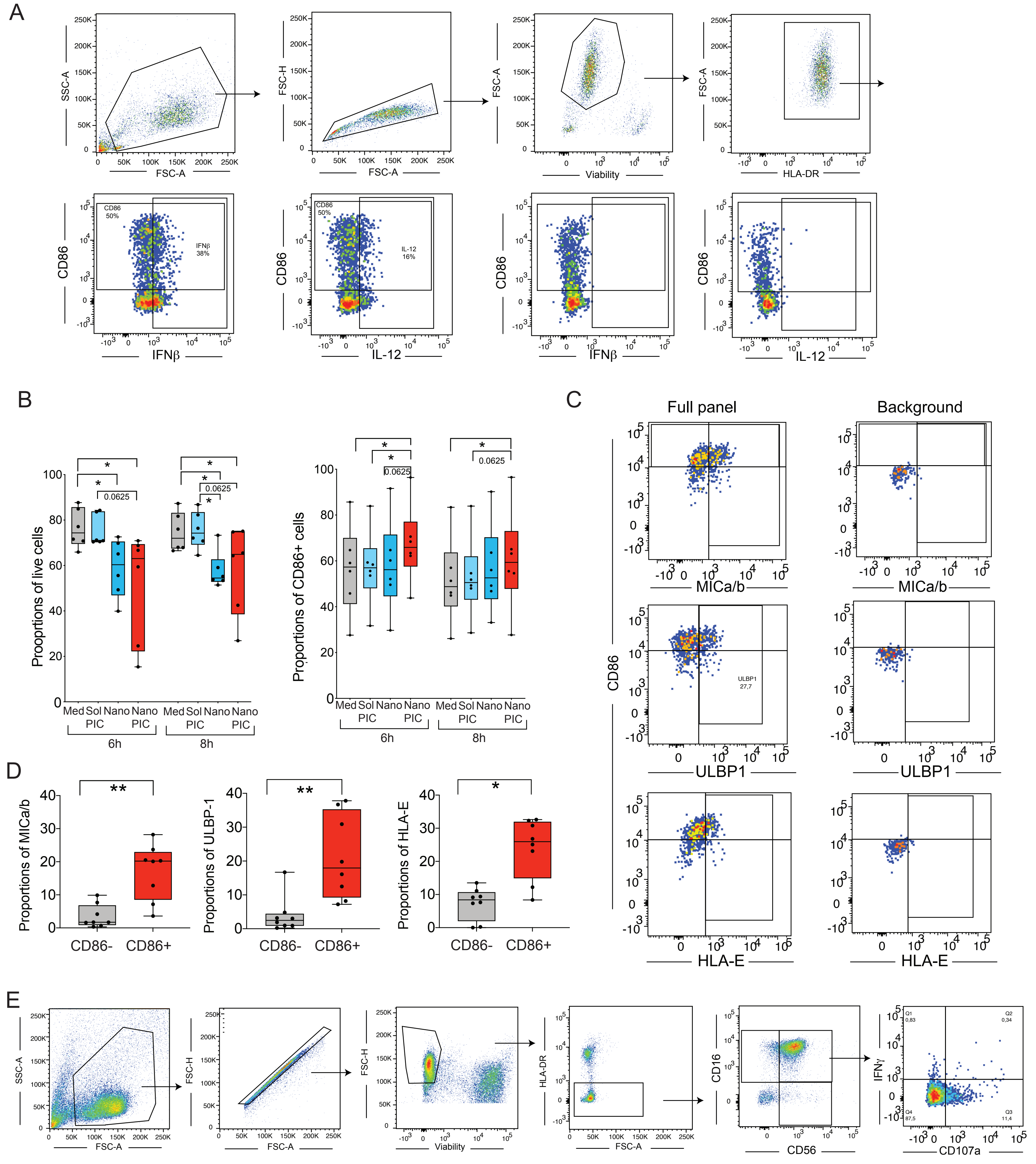

Supplemental Figure 2

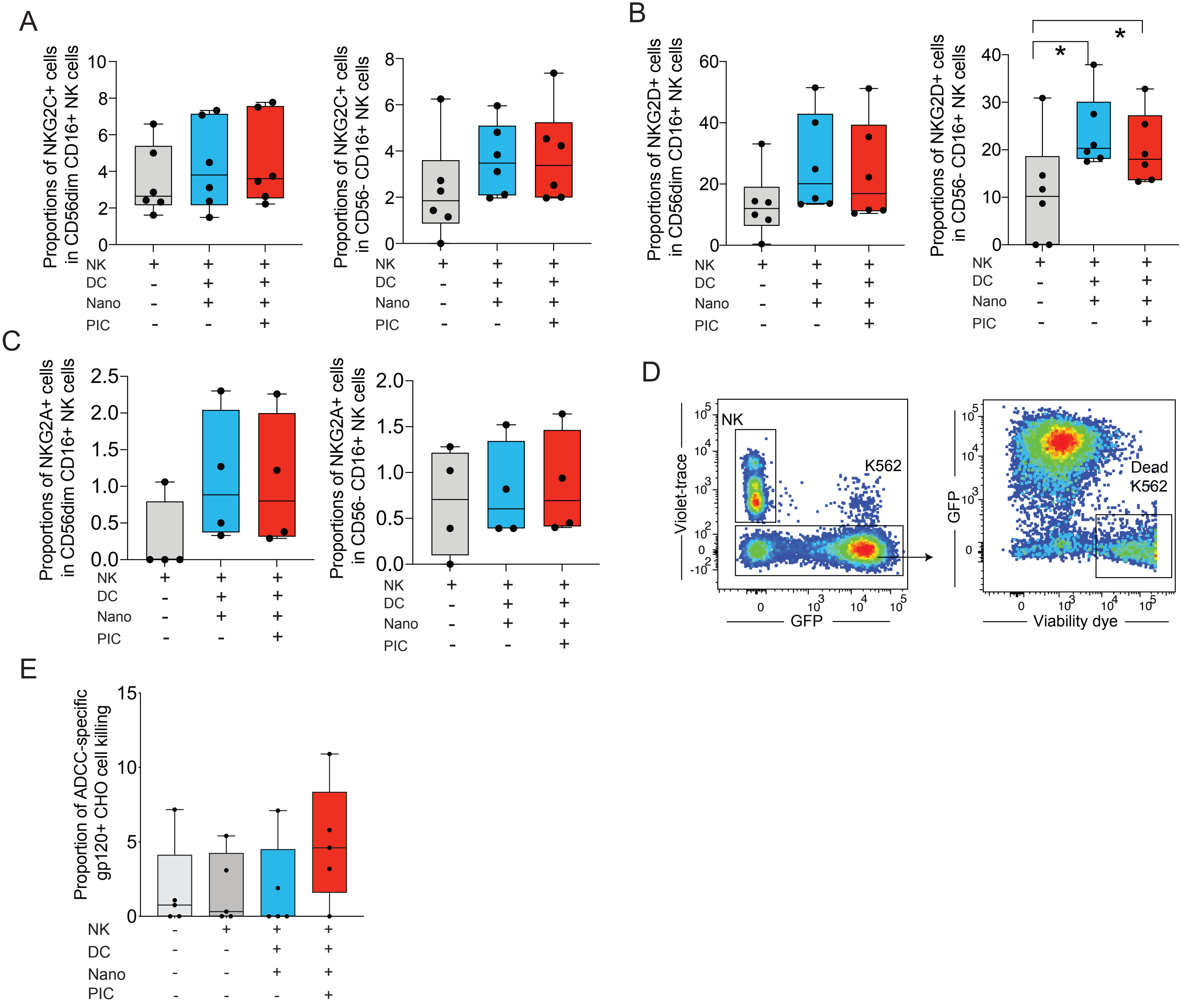

### Supplemental Figure 3

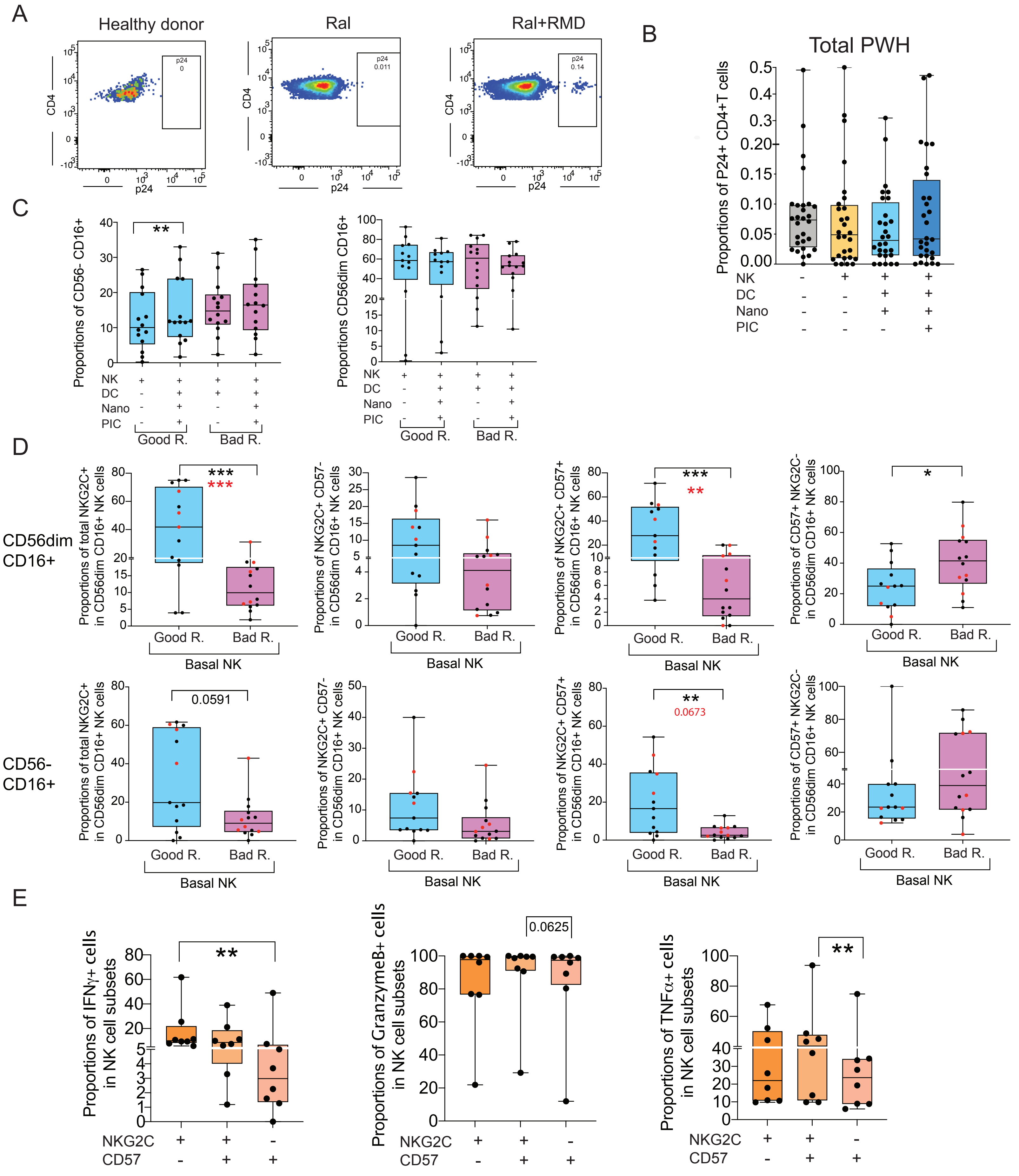

Supplemental Figure 4

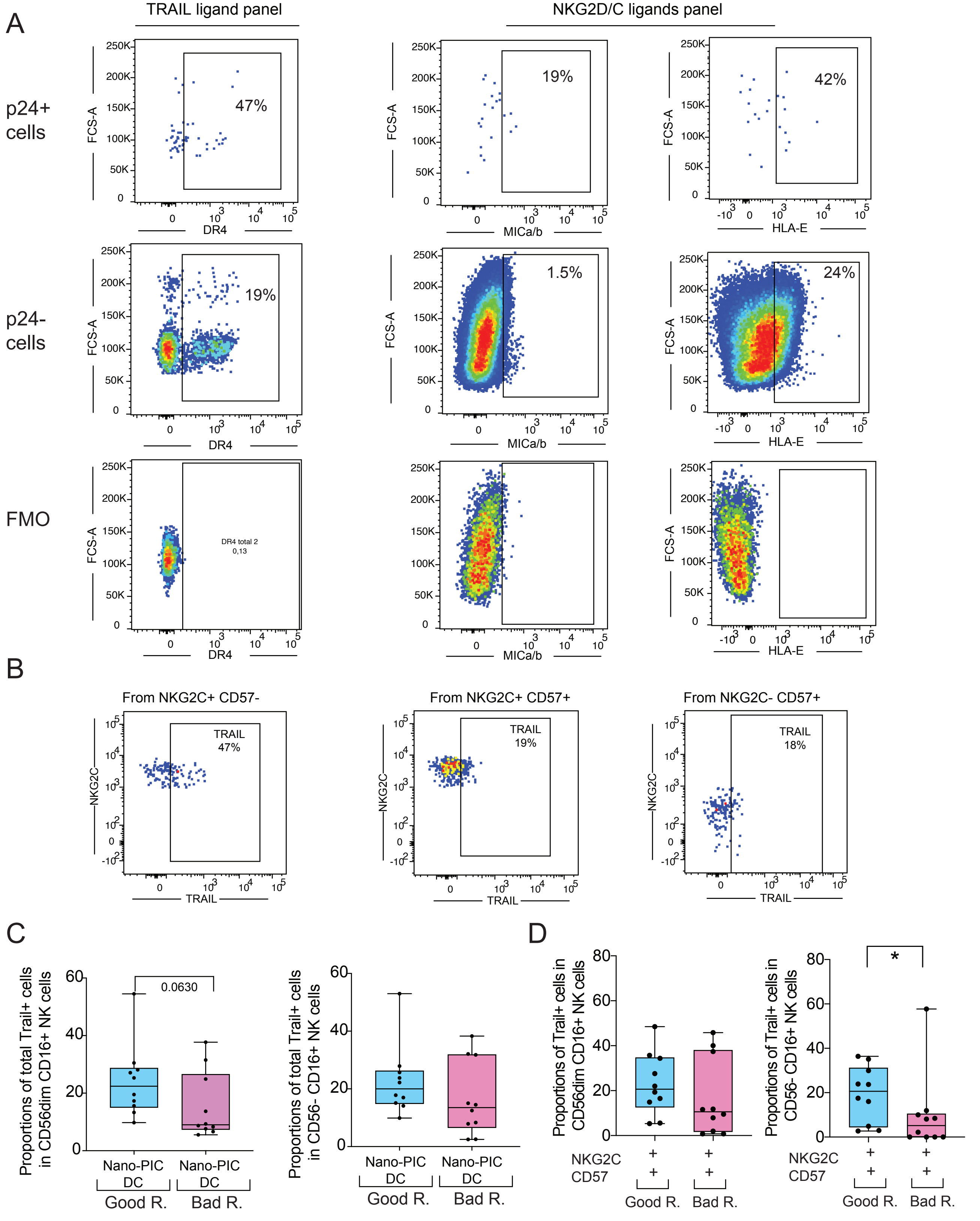

Supplemental Figure 5

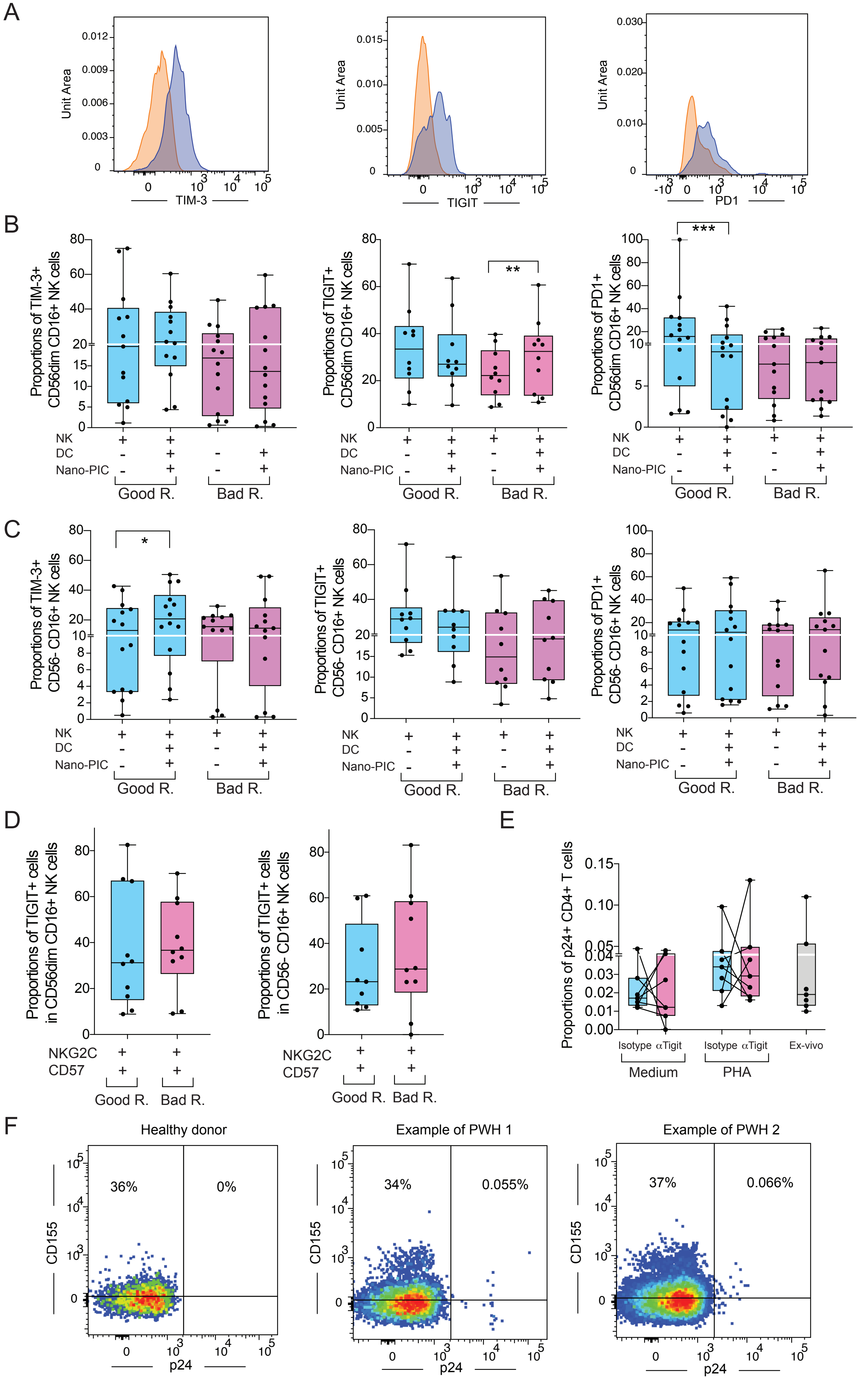

Supplemental Figure 6

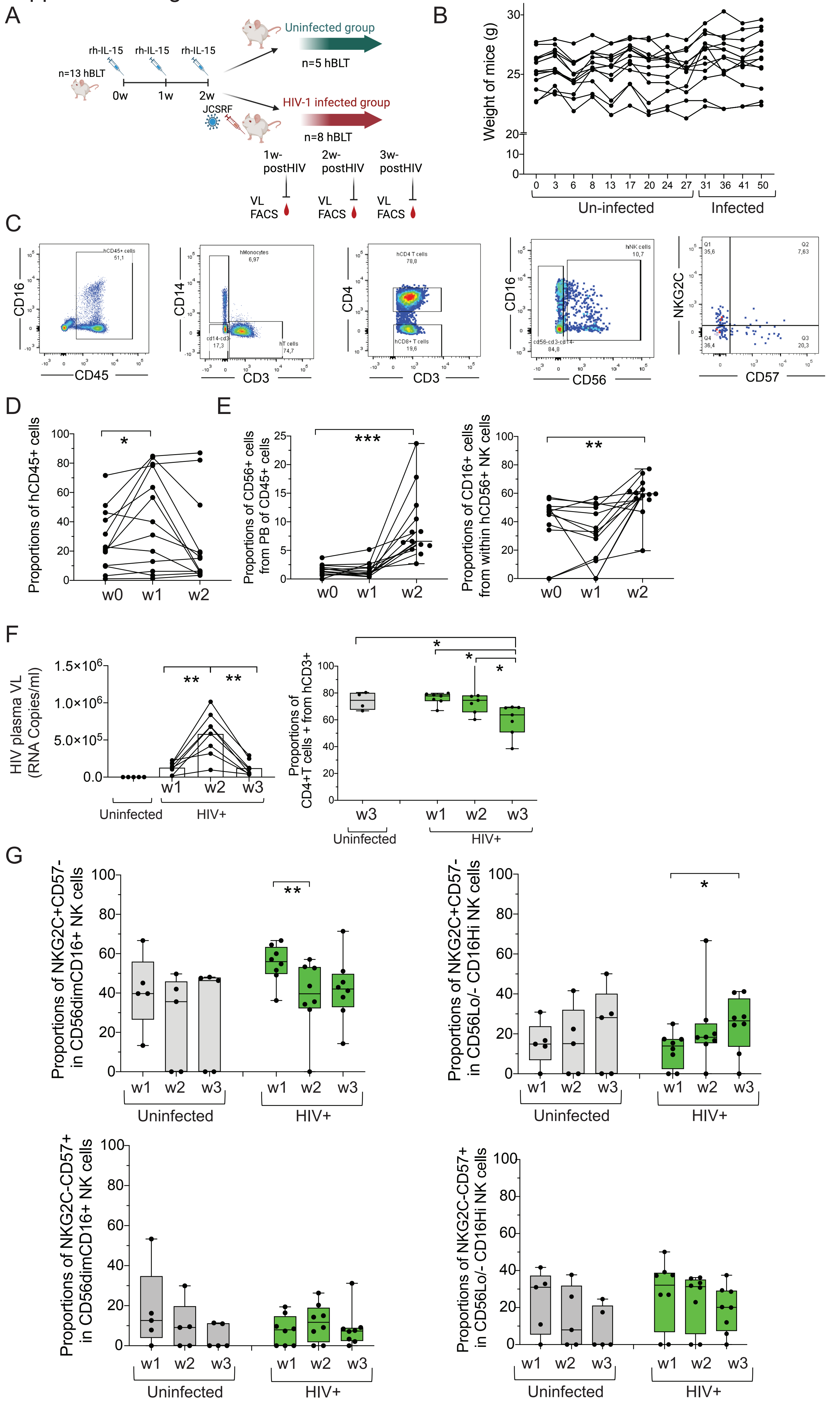

### Supplemental Figure 7

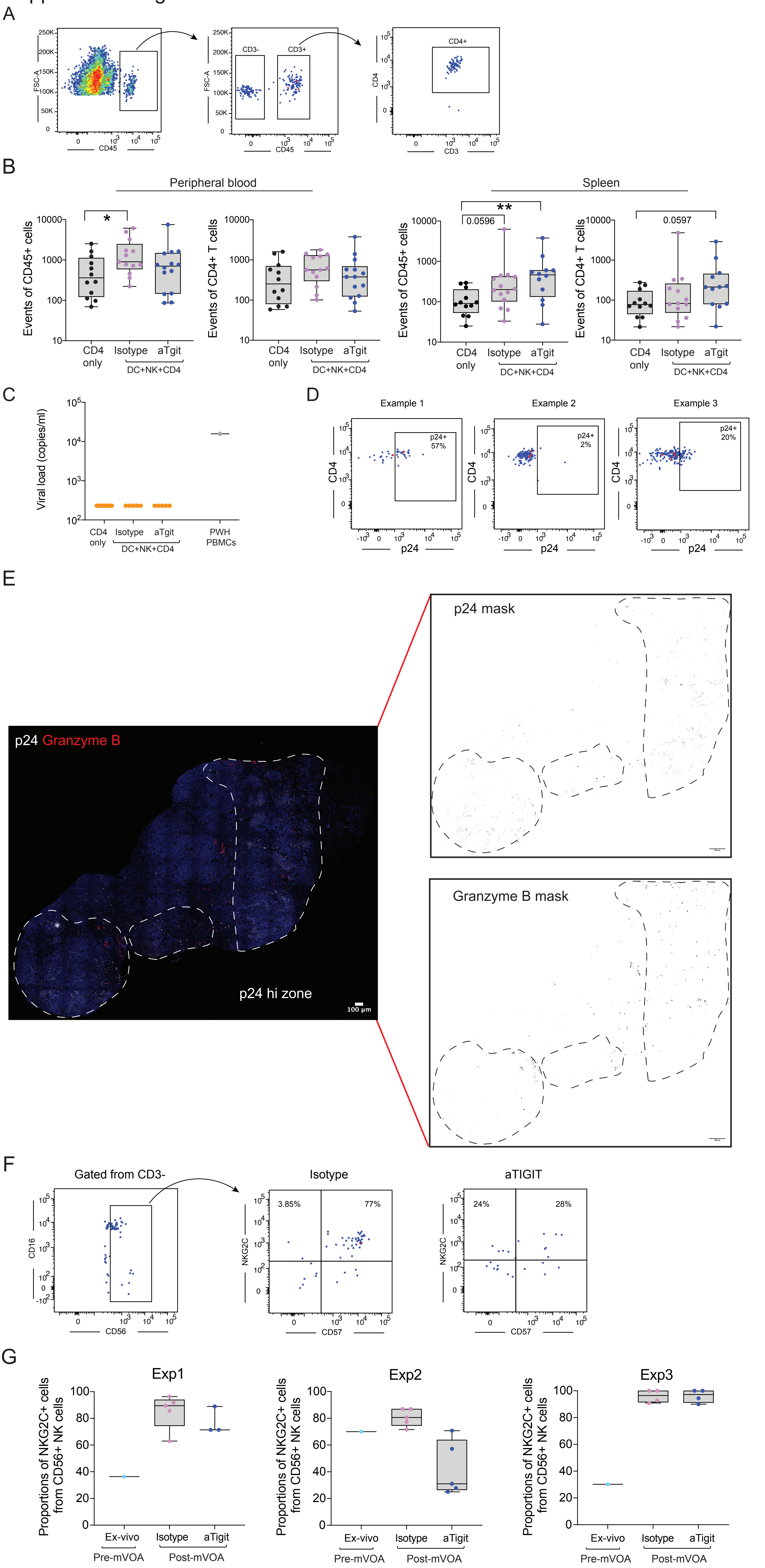

| | Total patients | Good-responders | Bad-responders | P values |
|---|---|---|---|---|
| **PWH n** | 28 | 14 | 14 | |
| **CD4 T cell count median (min-max)** | 835 (596-1639) | 969 (626-1639) | 829 (596-1586) | 0.34 |
| **CD4 T cells NADIR count median (min-max)** | 427 (9-869) | 452 (161-596) | 362 (9-869) | 0.25 |
| **Ratio CD4/CD8+ T cells count median (min-max)** | 1 (0.4-2.44) | 1.07 (0.51-2.44) | 0.89 (0.4-1.68) | 0.16 |
| **Age (years) median (min-max)** | 52 (25-75) | 44 (25-71) | 56 (26-75) | 0.01* |
| **Time on ART (Years) median (min-max)** | 9.2 (1.5-25) | 7 (1.5-25) | 13 (4 -24) | 0.11 |
| **Sex (% male)** | 96 | 100 | 91 | 0.82 |
| **Viral load at diagnosis (RNA copies/ml)** | 96000 (3810-2600000) | 96000 (3810-2600000) | 53000 (99-825000) | 0.48 |
| **CMV (% IgG positive)** | 70 | 78 | 62 | 0.07 |

**Supplemental Table 1. Clinical and demographic parameters of the PWH study cohort.**

#### Supplemental Table 2. Commercial reagents used for the study

| Reagent | Source | Catalog Number |
|---|---|---|
| **Cell lines** | | |
| EGFP-K562 | NIH Reagent Program | 116799 |
| HIV-1 HXB2 gp120 Expressing CHO cells (CHO-WT) | NIH Reagent Program | 2239 |
| HEK-293 | ATCC | CRL-1573 |
| **Antibodies** | | |
| Mouse Anti-TRAIL | RyD Systems | MAB375 |
| Mouse Anti-human NKG2C | RyD Systems | MAB1381 |
| Mouse Anti-human NKG2D | RyD Systems | MAB139 |
| IgG1 Isotype Control | BioLegend | 401402 |
| Mouse Anti-human TIGIT | RyD Systems | MAB7898 |
| Goat Anti-human TIM-3 | RyD Systems | AF2365 |
| IgG2b Isotype Control | BioLegend | 402202 |
| Purified Goat IgG | SIGMA | |
| Tiragolumab (anti-TIGIT) | Selleck Chemicals | A2028 |
| Human IgG1 isotype | BioXCell | BE0297 |
| Anti-human IFNβ | Pbl assay science | PBL-21400-3 #7225 |
| Anti-IL-12 | RyD Systems | IC2191C |
| Anti-human CD86 | Biolegend | 374210 |
| Anti-human ULBP1 | RyD systems | FAB1380P |
| Anti-human MICA/B | Biolegend | 320912 |
| Anti-human HLA-E | Biolegend | 342606 |
| Anti-human IFNγ | Biolegend | 554551 |
| Anti-human CD107a | Biolegend | 328620 |
| Anti-human TNFα | Biolegend | 502923 |
| Ghost Dye$^{TM}$ Red 780 | TONBO biosciences | 13-0865-T100 |
| Anti-human CD16 | Biolegend | 302045 |
| Anti-human CD56 | Biolegend | 362505 |
| Anti-human HLA-DR | Biolegend | 307630 |
| Anti-human PD-1 | Biolegend | 329938 |
| Anti-human CD253 (TRAIL) | Biolegend | 308210 |
| Anti-human DR4 (TRAIL-R1) | Biolegend | 307207 |
| Anti-human TIGIT (VSTM3) | Biolegend | 372734 |
| Anti-human CD45 | BD Horizon | 560777 |
| Anti-human CD155 (PVR) | Biolegend | 337614 |

| | | |
|---|---|---|
| Anti-human CD57 | Biolegend | 359624 |
| Anti-human NKG2A | RyD Systems | FAB1059C-025 |
| Anti-human NKG2D | Biolegend | 320819 |
| Anti-human NKG2C | RyD Systems | FAB138G |
| Anti-human NKG2C | RyD Systems | FAB138A |
| Anti-human CD3 | Biolegend | 300306 |
| Anti-human CD4 | Biolegend | 317429 |
| Anti-human P24 | Beckman Coulter | 6604667 |
| Anti-human Granzyme B | Biolegend | 563389 |
| Anti-human TIM3 | Biolegend | 345022 |
| Anti-human CD14 | Biolegend | 367110 |
| Rabbit anti-human NKG2C | Abcam | AB230900 |
| Goat anti-human TRAIL | RyD Systems | AF375 |
| Mouse anti-human p24 (Clone Kal-1) | Dako | M0857 |
| Granzyme B antibody | Thermo Fisher | 14-8889-80 |
| Donkey anti-rabbit AF488 | Thermo Fisher | R37118 |
| Donkey anti-rat AF594 | Jackson ImmunoResearch | 712-586-150 |
| Donkey anti-goat AF568 | Thermo Fisher | A-11057 |
| Donkey anti-mouse AF647 | Thermo Fisher | A-31571 |
| **Chemicals, Enzymes and other reagents** | | |
| Polyinosinic-polycytidylic acid sodium salt (poly I:C) | Merck- Sigma | P1530 |
| Brefeldin A | Sigma-Aldrich (Merck) | B7651 |
| Monensin | Sigma-Aldrich (Merck) | |
| Easy step human NK cell Isolation Kit | Stemcell | 17955 |
| PMA | Sigma Aldrich (Merck) | P1585 |
| Ionomicin | Sigma Aldrich (Merck) | I0634-1MG |
| Recombinant human IL-15 | RyD Systems | 247-ILB/CF |
| TransIT-X2 | Mirus Bio | MIR 6004 |
| **Software** | | |
| FlowJo v10 | | |
| Image J | | |
| Gradpad Prism 9.0 | | |
| **Other** | | |
| Illumina NextSeq | San Diego, CA | |
| Leica TCS SP5 confocal | Leica | |

| Good responders | | | | Bad responders | | | |
|---|---|---|---|---|---|---|---|
| Patients | Ral+RMD | Nano-PIC DC | Fold change | Patients | Ral+RMD | Nano-PIC DC | Fold change |
| **ART 24** | 0.18 | 0.034 | 0.18 | **ART 25** | 0.08 | 0.2 | 2.5 |
| **ART 22** | 0.014 | 0 | 0 | **ART 23** | 0.024 | 0.47 | 19.58 |
| **ART 30** | 0.49 | 0 | 0 | **ART 15** | 0.024 | 0.081 | 3.37 |
| **ART 43** | 0.09 | 0.001 | 0.01 | **ART 33** | 0.045 | 0.15 | 3.33 |
| **ART 40** | 0.28 | 0.023 | 0.08 | **ART 12** | 0.076 | 0.46 | 6.05 |
| **ART 31** | 0.078 | 0.028 | 0.35 | **ART 44** | 0.071 | 0.2 | 2.82 |
| **ART 8** | 0.098 | 0.069 | 0.7 | **ART 45** | 0.16 | 0.22 | 1.375 |
| **ART11** | 0.035 | 0 | 0 | **ART 29** | 0.1 | 0.1 | 1 |
| **ART20** | 0.026 | 0.01 | 0.38 | **ART 18** | 0.012 | 0.014 | 1.17 |
| **ART10** | 0.068 | 0.019 | 0.28 | **ART 7** | 0.021 | 0.037 | 1.76 |
| **ART50** | 0.038 | 0.00241 | 0.06 | **ART 54** | 0.042 | 0.11 | 2.62 |
| **ART55** | 0.096 | 0.041 | 0.43 | **ART 39** | 0 | 0.014 | 1.4 |
| **ART56** | 0.076 | 0.043 | 0.56 | **ART 26** | 0.1 | 0.11 | 1.1 |
| **ART17** | 0.098 | 0.087 | 0.89 | **ART 28** | 0.05 | 0.16 | 3.2 |

**Supplemental table 3. Change in proportions of p24+ CD4+ T cells from PWH after co-culture with NK and nano-PIC DC.** Data from individual donors included in the two good (blue) and bad (red) responder PWH groups are specified. Raw proportions of p24+ cells at baseline and after culture with NK and Nano-PIC DC are shown. Fold change in these proportions is also shown. Individuals that did not reach a fold change of <0.5 or >1.5 in p24+ frequencies are highlighted in red. Raltegavir (Ral); Romidepsin (RMD).

| | Donor exp 1 | Donor exp 2 | Donor exp 3 |
|---|---|---|---|
| **CD4 T cell count** | 1148 | 989 | 650 |
| **CD4 T cells NADIR count** | 420 | 404 | 172 |
| **Ratio CD4/CD8+ T cells count** | 0.78 | 1.2 | 1.1 |
| **Age (years)** | 62 | 40 | 48 |
| **Time on ART (Years)** | 16 | 9 | 9 |
| **Sex** | Male | Male | Male |
| **Viral load at diagnosis (RNA copies/ml)** | 53000 | 10063 | 183000 |
| **CMV IgG** | Positive | Positive | Positive |

**Supplemental Table 4. Clinical and demographic parameters of the PWH used in the 3 mVOA experiments.**
Experiment (Exp)
